# Developmental changes in the carotid body transcriptome accompanying the maturation of chemosensitivity

**DOI:** 10.1101/2025.08.21.671502

**Authors:** Emma J Hodson, Yoichiro Sugimoto, Rahul S Manamperige, Sage G Ford, Kimberley J Botting-Lawford, Youguo Niu, James A Nathan, Dino A Giussani, Peter J Ratcliffe

## Abstract

The carotid body (CB) chemoreceptors mediate rapid cardiorespiratory reflexes to hypoxia, which mature peri-natally and are vital for fetal hypoxia tolerance and post-natal ventilatory control. This maturation is associated with an increase in the sensitivity of the CB electrophysiological response to hypoxia (chemosensitivity): a process that is incompletely understood but critical to systemic oxygen homeostasis. Hypothesizing that perinatal CB gene expression changes would reveal candidate mechanisms for oxygen chemosensitivity, we studied the CB transcriptome in sheep, where peri-natal CB physiology is well-characterised. CB-mediated cardiovascular reflexes are detectable at fetal day 120, and robust by term (day 145), while hypoxic ventilatory responses are established by post-natal day 15. We performed RNA sequencing on sheep CBs at each of these stages, and adults, along with the superior cervical ganglion (SCG) as an oxygen-insensitive control. This allowed us to define tissue-specific changes in the CB transcriptome correlating with chemosensitivity maturation. Striking, progressive CB enrichment is observed in genes implicated in murine CB chemosensitivity, including potassium channels (*KCNK9*), mitochondrial complex IV regulators (*NDUFA4L2, HIGD1C*), and HIF-2α (*EPAS1*). Genes with this expression pattern are also enriched for regulators of diacylglycerol (DAG), particularly the DAG kinase *DGKH:* one of the most abundant CB transcripts increasing in parallel with chemosensitivity. Across developmental stages, the CB also exhibits marked down-regulation of metabolic pathways and ATP/GTP consuming processes, potentially providing a state permissive to metabolic signal detection. Together, this builds a detailed picture of the CB transcriptional signature, with core features established in fetal life and conserved across species.

**Key points:**

- The carotid body (CB) chemoreceptors mediate rapid cardiorespiratory responses to hypoxia, which maintain systemic oxygen homeostasis, but CB dysfunction is also implicated in pathologies including hypertension, heart failure and sudden infant death.
- CB-mediated chemoreflexes mature during the peri-natal period, with increasing sensitivity of the oxygen chemosensory response.
- We performed RNA-seq of CBs from sheep across 3 peri-natal stages and adults, enabling us to identify gene expression changes that correlate with functional state.
- We describe a CB transcriptomic signature that is conserved across species, established in fetal life, and correlates with maturation. This includes features of a unique metabolic phenotype, and up-regulation of genes encoding the extracellular matrix and diacylglycerol signalling. The top transcription factor correlating with functional maturation is *EPAS1*/ HIF-2α.
- We anticipate that this data set will be a valuable resource in generating novel hypotheses on mechanisms of oxygen chemosensory function, development and CB-associated pathologies.

## Introduction

The carotid body arterial chemoreceptors play a crucial role in systemic oxygen homeostasis, mediating rapid cardiorespiratory reflexes in response to hypoxia that serve to maintain arterial oxygen levels (Kumar & Prabhakar, 2012). Chemoreceptor reflexes undergo complex developmental changes during the peri-natal period with the transition to airbreathing life and the associated increase in oxygen uptake and delivery. During this time, carotid body (CB) function is essential both for fetal hypoxia tolerance, through regulation of a cardiovascular chemoreflex (Giussani *et al*., 1993; Giussani, 2016), and for establishing stable ventilatory control in the neonate (Carroll, 2002; Carroll & Agarwal, 2010). While being critical in the maintenance of physiological defences against acute hypoxaemia during the peri-natal period, the CB must simultaneously adapt to the dramatic increase in oxygen availability at birth by adjusting the chemosensory activation setpoint to match the newly elevated normalcy for arterial oxygen. This involves an increase in the sensitivity of the chemosensory response to hypoxia, which has been demonstrated at the level of cardiorespiratory reflexes, CB afferent nerve activity, and electrophysiological responses of individual CB cells (reviewed in Carroll & Kim, 2013). Understanding these important adaptations in chemosensory function may provide insights into the pathophysiology of stillbirth, neonatal apnoea and sudden infant death syndrome, which are associated with peri-natal hypoxia and CB pathology (MacFarlane *et al*., 2013; Neary & Breckenridge, 2013; Porzionato *et al*., 2018; Pacora *et al*., 2019; Lear *et al*., 2024). Studying this developmental functional maturation may also shed light on normal CB physiology and the mechanisms underlying oxygen chemosensitivity, which remain incompletely understood.

CB chemosensory cells are neuroendocrine cells, known as type I or glomus cells, which exhibit a rapid neurosecretory response to hypoxia. This is proposed to involve depolarisation of the plasma membrane through closure of potassium channels (TASK1/3, maxi-K), and activation of voltage-gated calcium channels with the resulting calcium influx triggering neurosecretion (López-Barneo *et al*., 1988; Buckler & Vaughan-Jones, 1994; Buckler, 1997, reviewed in López-Barneo *et al*., 2016). The mitochondria also appear to be important for the chemosensory response as inhibitors of the electron transport chain are potent stimulators of CB type I cell activity (Mulligan *et al*., 1981; Mulligan & Lahiri, 1982; Wyatt & Buckler, 2004; Turner & Buckler, 2013), and type I cell mitochondria, specifically mitochondrial complex IV, are particularly sensitive to moderate hypoxia (Mills & Jöbsis, 1970, 1972; Nair *et al*., 1986; Duchen & Biscoe, 1992a, 1992b; Buckler & Turner, 2013).

Studies of CB gene expression have been highly informative in developing our understanding of CB type I cell biology and have highlighted overexpression of certain unusual isoforms of genes contributing to the function of complex IV (*COX4I2, NDUF4AL2, HIGD1C*) and the vertebrate-specific isoform of Hypoxia-inducible factor alpha (*EPAS1*/ HIF-2α) (Chang *et al*., 2015; Zhou *et al*., 2016; Gao *et al*., 2017). The high absolute and relative expression levels of these genes in CB tissue suggests a causative role in oxygen chemosensitivity, which in some cases is supported by functional studies with genetic or pharmacological interventions (Hodson *et al*., 2016; Fielding *et al*., 2018; Cheng *et al*., 2020; Moreno-Domínguez *et al*., 2020; Timón-Gómez *et al*., 2022; Prange-Barczynska *et al*., 2023, reviewed in Gao *et al*., 2025). However, there are outstanding questions with respect to how changes in oxygen concentration are detected and how the signal is communicated to ion channels at the plasma membrane. There are also some limitations to the existing literature, the large majority of which concerns small rodents, so may repeatedly highlight species-specific gene expression patterns.

To focus gene expression studies on those most likely to have a fundamental role in CB function, we have studied the change in CB gene expression from fetal to adult life, across a period corresponding to an increase in oxygen chemosensitivity in experimental animals studied. We have used the sheep as a physiological model, as it offers several substantive advantages. First, sheep and humans share similar milestones in fetal cardiovascular development, therefore fetal studies using sheep are of high human translational value (Morrison *et al*., 2018). Second, there is comprehensive background physiological data on the maturation of CB functions during the progression from fetal to adult life in sheep, allowing us to compare different developmental stages that have been well characterised with respect to oxygen chemosensitivity. Third, the ovine CB is much larger than that of the small rodents used in most studies to date, enabling high depth RNA-sequencing analyses to be conducted on individual CB organs (as opposed to pooled samples), even at fetal stages, which have not been included in previous RNA-sequencing studies. Finally, using the sheep enabled us to refine the list of genes showing conserved association with chemosensitivity across species.

To facilitate comparison of gene expression data with functional studies we chose to analyse the following four time points (Figure 1A):

a. Fetal day 120, which is the earliest stage when CB-mediated reflexes are well characterised, with hypoxic stimulation leading to bradycardia and peripheral vasoconstriction, part of the fetal brain sparing circulatory response to acute hypoxia (Bartelds *et al*., 1993; Giussani *et al*., 1993, 2001);
b. Fetal day 145, which is full term in the breed of sheep we used (Brain *et al*., 2019). Near term, the CB-mediated fetal brain sparing circulatory to hypoxia response becomes more robust; increasing in magnitude (Fletcher *et al*., 2006). Recordings of CB afferent nerve activity suggest that this is, at least in part, due to intrinsic gains in CB oxygen sensitivity (Blanco *et al*., 1984);
c. Post-natal day 15 lambs, when stable respiratory control is established with robust CB-mediated hypoxic ventilatory responses. Prior to this, the immediate neonatal period is associated with respiratory instability and poor hypoxic ventilatory responses, principally due to a weak CB chemoreceptor response (Bureau & Begin, 1982; Mayock *et al*., 1983; Blanco *et al*., 1984; Bureau *et al*., 1985b, 1985a; Canet *et al*., 1996). At birth, the CB is relatively quiescent, with a threshold for activation still corresponding to the hypoxic fetal circulation (Blanco *et al*., 1984). As arterial oxygen levels rise following transition from placental to pulmonary gas transfer, CB activation shifts to a higher oxygen setpoint during the first 2 weeks of post-natal life. This involves a CB cell intrinsic gain in oxygen sensitivity; as shown by calcium and potassium channel responses in isolated type I cells, and carotid sinus nerve recordings (Bamford *et al*., 1999; Wasicko *et al*., 1999, 2006; Kim *et al*., 2011);
d. Adult sheep, when post-natal growth and CB maturation are complete.

**Figure 1.**
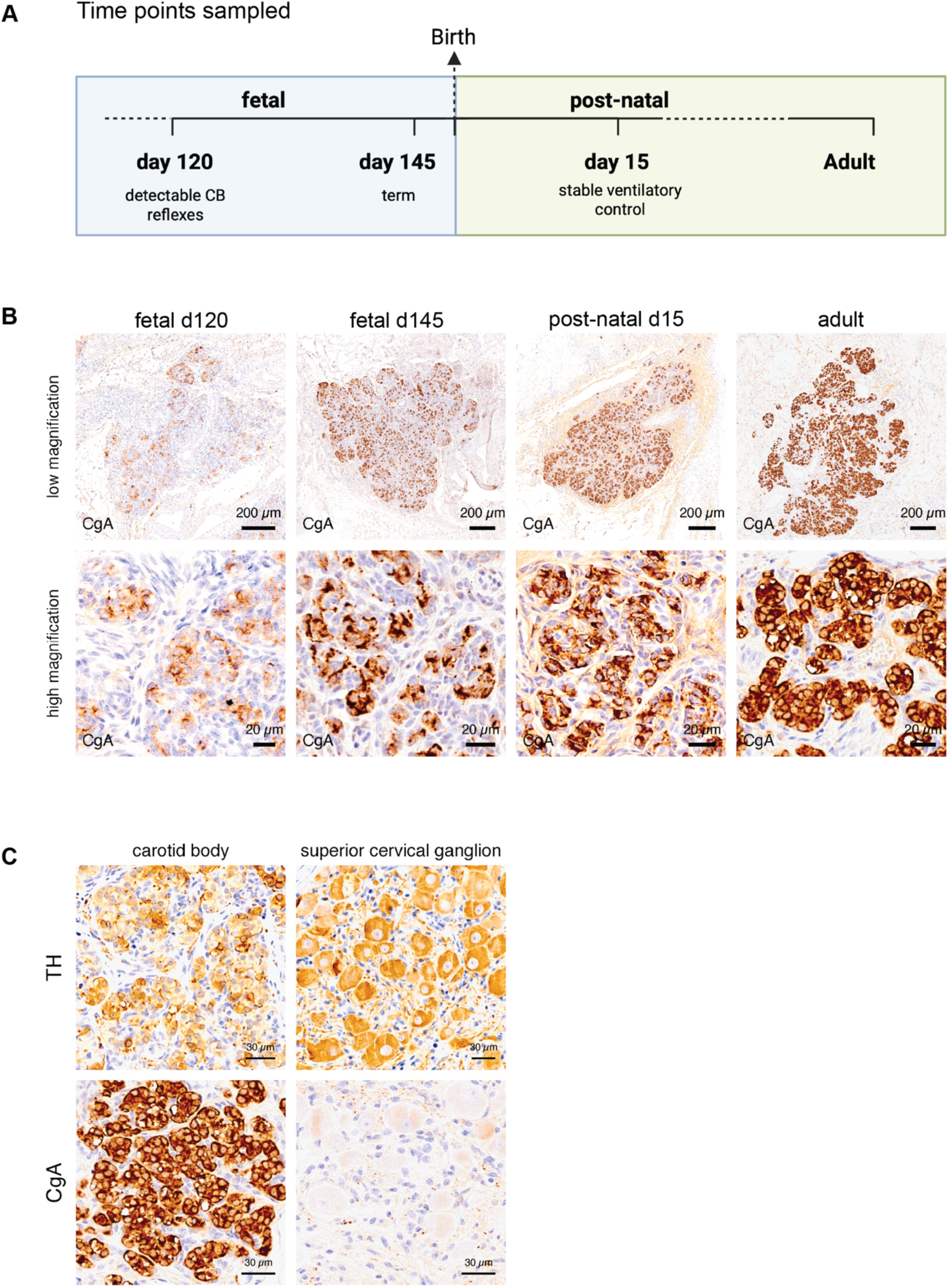
Experimental outline and identification of carotid body and superior cervical ganglion. (A) Timeline of developmental stages sampled. (B) Immunostaining for chromogranin A (CgA) on sheep CB from each developmental stage shown in (A). (C) Immunostaining for tyrosine hydroxylase (TH) and chromogranin A (CgA) on carotid body (CB) and super cervical ganglion (SCG) tissues from adult sheep. Scale bar= 200 uM (upper panels) and 20 uM (lower panels).

In parallel, we wished to compare CB gene expression changes to those in a tissue that does not acquire oxygen chemo-sensitivity and hence focus on genes that show both tissue-specific and developmentally regulated changes in expression. We chose the superior cervical (sympathetic) ganglion (SCG), which has physiological and ontological features in common with the CB but does not manifest oxygen chemosensitivity. The SCG has been used as a control tissue in previous studies of CB gene expression (Chang *et al*., 2015; Gao *et al*., 2017), facilitating the integration of our data with that work.

## Results

The sheep CB is found at the origin of the occipital artery and can be readily identified as a defined golden oval/lobular structure at the intersection of the artery with the ganglioglomerular and carotid sinus nerves, which respectively form the efferent and afferent CB innervation. CB identity was confirmed by histology, showing the classic nested “zellballen” morphology of type I cells staining positive for tyrosine hydroxylase (TH) and chromogranin A, with signal intensity increasing with maturation (Figure 1B, C). The SCG is located at the opposite end of the (sympathetic) ganglioglomerular nerve and comprises sympathetic neuronal cell bodies that stain positive for TH, as these cells also produce catecholamines, but not the neuroendocrine marker chromogranin A (Figure 1C).

RNA was extracted from three samples from each tissue and time point for RNA sequencing. One SCG sample (d145) was excluded due to poor RNA quality and limited expression of sympathetic markers. All other samples were included in the analysis, and we provide this dataset in full as a resource available on Github (see *Methods*).

### Comparative analysis of carotid body-specific gene expression

We first sought to define genes that show the most striking contrast in expression levels between the sheep CB and SCG, with an initial focus on samples from the adult stage, where oxygen chemosensitivity is maximally manifest, and fetal day 120, which is developmentally the most distinct from the adult data, and novel as there are no previous studies of this type incorporating the fetal CB transcriptome.

Our analysis identified 9,335 genes in the adult and 6,555 genes from fetal day 120 with significant differential expression between the CB and SCG (P<0.01). Genes showing the greatest fold change either up (CB-enriched) or down-regulated (SCG-enriched) are shown in Table 1A-D.

**Table 1:** Genes with the greatest differential expression between CB and SCG in the adult and fetal d120 sheep. (A-B) The top 20 genes most up-regulated (A) and down-regulated (B) in the adult sheep CB versus SCG. (C-D) The top 20 genes most up-regulated (C) and down-regulated (D) in the fetal day 120 sheep CB versus SCG. Genes are ranked by log2 fold change (log2FC). *P value less than 1E-20. All values shown to two significant figures.

| Gene name | Description | log2FC | P value |
| --- | --- | --- | --- |
| FGF19 | Fibroblast growth factor 19 | 15.2 | * |
| INSM1 | Insulinoma-Associated Protein 1 | 14.2 | * |
| HIGD1C | HIG1 Hypoxia Inducible Domain Family Member 1C | 13.2 | * |
| GPR139 | G Protein-Coupled Receptor 139 | 11.6 | * |
| NEUROD4 | Neuronal Differentiation 4 | 11.4 | 2.6E-18 |
| EPYC | Epiphycan | 11.3 | 2.5E-09 |
| HMX3 | H6 Family Homeobox 3 | 10.4 | * |
| DLX6 | Distal-Less Homeobox 6 | 10.2 | * |
| KCNK9 | Potassium Two Pore Domain Channel Subfamily K Member 9 | 10.2 | * |
| ACSM1 | Acyl-CoA Synthetase Medium Chain Family Member 1 | 9.9 | 2.3E-14 |
| NTS | Neurotensin | 9.9 | 2.9E-14 |
| SLC26A4 | Pendrin (Chloride transporter) | 9.5 | * |
| GRIN2B | Glutamate Ionotropic Receptor NMDA Type Subunit 2B | 9.5 | 1.8E-15 |
| NXPH4 | Neurexophilin 4 | 9.5 | * |
| GLRA1 | Glycine Receptor Alpha 1 | 9.3 | * |
| NOTUM | Notum, Palmitoleoyl-Protein Carboxylesterase | 9.1 | * |
| RSPO1 | R-Spondin 1 | 8.9 | 1.8E-11 |
| HMX2 | H6 Family Homeobox 2 | 8.6 | 2.4E-14 |
| B3GAT2 | Beta-1,3-Glucuronyltransferase 2 | 8.6 | * |
| CA4 | Carbonic Anhydrase 4 | 8.5 | 4.5E-18 |

| Gene name | Description | log2FC | P value |
| --- | --- | --- | --- |
| GPX2 | Glutathione Peroxidase 2 | -16 | * |
| ECEL1 | Endothelin Converting Enzyme Like 1 | -12 | 3.1E-15 |
| HMX1 | H6 Family Homeobox 1 | -12 | * |
| NTRK1 | Neurotrophic Receptor Tyrosine Kinase 1 | -11 | * |
| CHRNA4 | Cholinergic Receptor Nicotinic Alpha 4 Subunit | -11 | 1.0E-18 |
| GRP | Gastrin Releasing Peptide | -11 | * |
| HTR3A | 5-Hydroxytryptamine Receptor 3A | -10 | 1.3E-11 |
| CPA6 | Carboxypeptidase A6 | -10 | 2.5E-16 |
| FSTL5 | Follistatin Like 5 | -10 | 1.1E-06 |
| DRGX | Dorsal Root Ganglia Homeobox | -10 | 1.6E-13 |
| AMER3 | APC Membrane Recruitment Protein 3 | -9.8 | * |
| PDZK1IP1 | PDZK1 Interacting Protein 1 | -9.7 | 7.0E-20 |
| DBH | Dopamine Beta-Hydroxylase | -9.5 | * |
| CALB2 | Calbindin 2 | -9.5 | * |
| SLC51B | SLC51 Subunit Beta | -9.4 | 2.4E-12 |
| RHBG | Rh Family B Glycoprotein | -9.3 | 3.1E-15 |
| HTR3B | 5-Hydroxytryptamine Receptor 3B | -9.3 | 1.8E-10 |
| CACNG8 | Calcium Voltage-Gated Channel Auxiliary Subunit Gamma 8 | -9.2 | 3.8E-10 |
| CRHR2 | Corticotropin Releasing Hormone Receptor 2 | -9.1 | * |
| IGSF5 | Immunoglobulin Superfamily Member 5 | -8.9 | 8.0E-12 |

| Gene name | Description | log2FC | P value |
| --- | --- | --- | --- |
| GPR139 | G Protein-Coupled Receptor 139 | 13 | * |
| FGF19 | Fibroblast growth factor 19 | 13 | * |
| NTS | Neurotensin | 12 | 5.4E-20 |
| LOC105603737 | uncharacterised | 10 | 1.9E-06 |
| HMX2 | H6 Family Homeobox 2 | 10 | 2.1E-15 |
| DLX6 | Distal-Less Homeobox 6 | 10 | * |
| CATHL1 | Cathelicidin-1 | 9.9 | 8.2E-04 |
| AGTR2 | Angiotensin II Receptor Type 2 | 9.8 | 5.0E-12 |
| CA4 | Carbonic Anhydrase 4 | 9.7 | * |
| HIGD1C | HIG1 Hypoxia Inducible Domain Family Member 1C | 9.3 | 1.1E-15 |
| HMX3 | H6 Family Homeobox 3 | 9.2 | * |
| TPH1 | Tryptophan Hydroxylase 1 | 9.1 | * |
| NOTUM | Notum, Palmitoleoyl-Protein Carboxylesterase | 9.0 | * |
| ACSM1 | Acyl-CoA Synthetase Medium Chain Family Member 1 | 8.8 | 2.8E-08 |
| CPS1 | Carbamoyl-Phosphate Synthase 1 | 8.8 | * |
| IGFBP1 | Insulin Like Growth Factor Binding Protein 1 | 8.7 | 9.7E-09 |
| NMUR2 | Neuromedin U Receptor 2 | 8.5 | 1.7E-11 |
| GJD2 | Gap Junction Protein Delta 2 | 8.4 | 3.1E-09 |
| CATHL3 | Cathelicidin-3 | 8.4 | 6.5E-07 |
| NXPH4 | Neurexophilin 4 | 8.2 | * |

| Gene name | Description | Log2FC | P Value |
| --- | --- | --- | --- |
| GPX2 | Glutathione Peroxidase 2 | -12 | 1.4E-21 |
| ECEL1 | Endothelin Converting Enzyme Like 1 | -11 | * |
| NTRK1 | Neurotrophic Receptor Tyrosine Kinase 1 | -11 | * |
| CHRNA4 | Cholinergic Receptor Nicotinic Alpha 4 Subunit | -9.4 | * |
| NAA80 | N-Alpha-Acetyltransferase 80, NatH Catalytic Subunit | -9.2 | 7.1E-12 |
| CCDC105 | Tektin-like protein 1 | -9.0 | 1.1E-11 |
| PKD1L2 | Polycystin-1-like protein 2 | -8.9 | * |
| TMEM275 | Transmembrane protein 275 | -8.9 | 3.0E-11 |
| NTSR1 | Neurotensin receptor type 1 | -8.8 | 1.1E-19 |
| HEPHL1 | Hephaestin-like protein 1 | -8.8 | 3.3E-11 |
| PRSS56 | Serine protease 56 | -8.6 | 6.5E-11 |
| HMX1 | Homeobox protein HMX1 | -8.4 | 1.8E-11 |
| DBH | Dopamine Beta-Hydroxylase | -8.3 | * |
| LOC101119944 | elongation factor 1-alpha 2-like | -8.2 | 3.0E-09 |
| TRHR | Thyrotropin-releasing hormone receptor | -8.1 | 4.8E-08 |
| NPSR1 | Neuropeptide S receptor | -8.0 | 7.6E-08 |
| SGPP2 | Sphingosine-1-phosphate phosphatase 2 | -8.0 | 2.4E-15 |
| SERTM2 | Serine-rich and transmembrane domain-containing 2 | -7.9 | * |
| SLC6A2 | Norepinephrine Transporter | -7.8 | * |
| TM4SF4 | Transmembrane 4 L6 family member 4 | -7.7 | 3.3E-14 |

Several previous studies, principally in mice, have examined CB gene expression with respect to control tissues (Chang *et al*., 2015; Zhou *et al*., 2016; Gao *et al*., 2017) and the present study provides a parallel dataset in a species distinct from small rodents. Genes that show tissue-specific expression patterns conserved between species are likely to be enriched for those with critical functional roles, including chemosensitivity. We therefore examined the concordance between the most highly CB-enriched genes identified in the sheep and those defined in earlier studies on small rodents. To collate comparable mouse datasets, we identified published studies describing transcriptomic data in the mouse CB with respect to at least one control tissue. We combined data from an RNA sequencing analysis comparing mouse CB and adrenal medulla (Chang *et al*., 2015); a microarray analysis comparing mouse CB and SCG (Gao *et al*., 2017); and a single cell RNAseq analysis of mouse CB type I cells compared to a range of control cell types (Zhou *et al*., 2016). Combining the lists of top-ranked CB-enriched genes in each study, we compiled a list of genes highly enriched in the mouse CB (n=168, Figure 2A-a-d) and used this to form a qualitative comparison with the top 160 most up-regulated genes in the adult (Figure 2A-b-d-e-g) and fetal sheep CB (Figure 2A-c-d-e-f).

**Figure 2.**
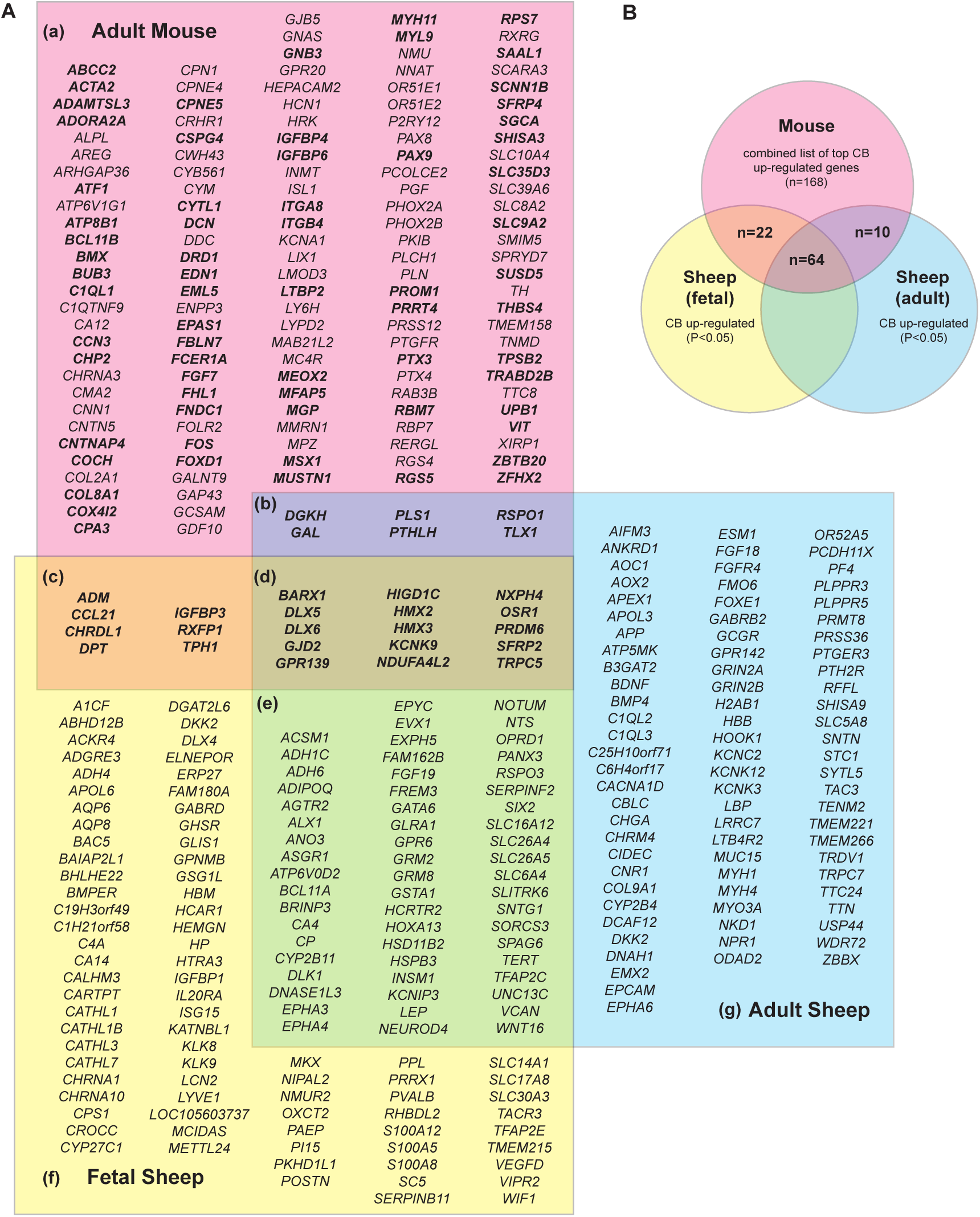
Overlap between the most up-regulated genes in mouse and sheep reveals a conserved CB gene expression signature. (A) Overlap between lists of the top-ranked genes up-regulated in the mouse or sheep CB: a-d (pink box) compiled list of the most up-regulated genes in mouse CB, relative to various control tissues (see text); b,d,e,g (blue box) the top 160 most up-regulated genes in the adult sheep CB versus SCG; c,d,e,f (yellow box) the top 160 most upregulated genes in the fetal d120 sheep CB versus SCG. (b-d) shows genes common to the mouse list and the fetal and/or adult sheep list. (B) Venn diagram showing the number of genes in the combined mouse list that are also significantly up-regulated in the adult and/or fetal sheep CB (total n=96). These genes are highlighted in bold in (A).

The top-ranked gene lists from the different mouse studies showed relatively modest overlap, perhaps reflecting the different control tissues used. However, the sheep CB gene list showed good concordance with the combined mouse data, with 28 genes common to mouse and either of the sheep top gene lists (Figure 2A-b-c-d). Of these, 15 were top hits in the mouse, and both adult and fetal sheep (Figure 2A-d). Extending the analysis to any gene with significant up-regulation in the sheep CB vs SCG (P<0.05) extended the overlap: 77/168 mouse CB genes were significantly upregulated in the adult sheep CB, and 86/168 were significantly upregulated in the fetal sheep CB, with 96/168 mouse genes being significantly upregulated in at least one sheep dataset (Figure 2B and 2A-a-d – genes in bold).

Volcano plots of differential gene expression between sheep CB and SCG in adults (Figure 3B) and the day 120 fetus (Figure 3A) illustrate the genes with the most striking CB-enriched expression, highlighting those that are also among the most up-regulated genes in the mouse CB (Figure 3, pink annotation) and others that are among the most significantly up-regulated transcripts in the sheep CB, but were not highlighted in mouse studies (Figure 3, blue annotation). The common gene expression profile, detected in both species, includes genes with proposed roles in oxygen chemosensitivity (eg *HIGD1C, NDUFA4L2, COX4I2, EPAS1/HIF-2, KCNK9*), and genes highly expressed in other tissues that appear to support oxygen chemosensitivity-like responses (*RGS5, DGKH*) (Hanemaaijer *et al*., 2021; Akkuratova *et al*., 2022). The CB-specific expression of these genes that we independently identify in sheep further enhances the evidence for functional relevance of these genes, which together form a CB gene expression signature that is established in fetal life. In contrast, there are several genes highlighted in mouse studies that we do not find to be upregulated in the sheep CB, including *Olfr78/OR51E2* and other olfactory receptors, for which some, but not all studies have proposed a functional role in the mouse CB (Chang *et al*., 2015; Torres-Torrelo *et al*., 2018; Peng *et al*., 2020; Colinas *et al*., 2024).

**Figure 3.**
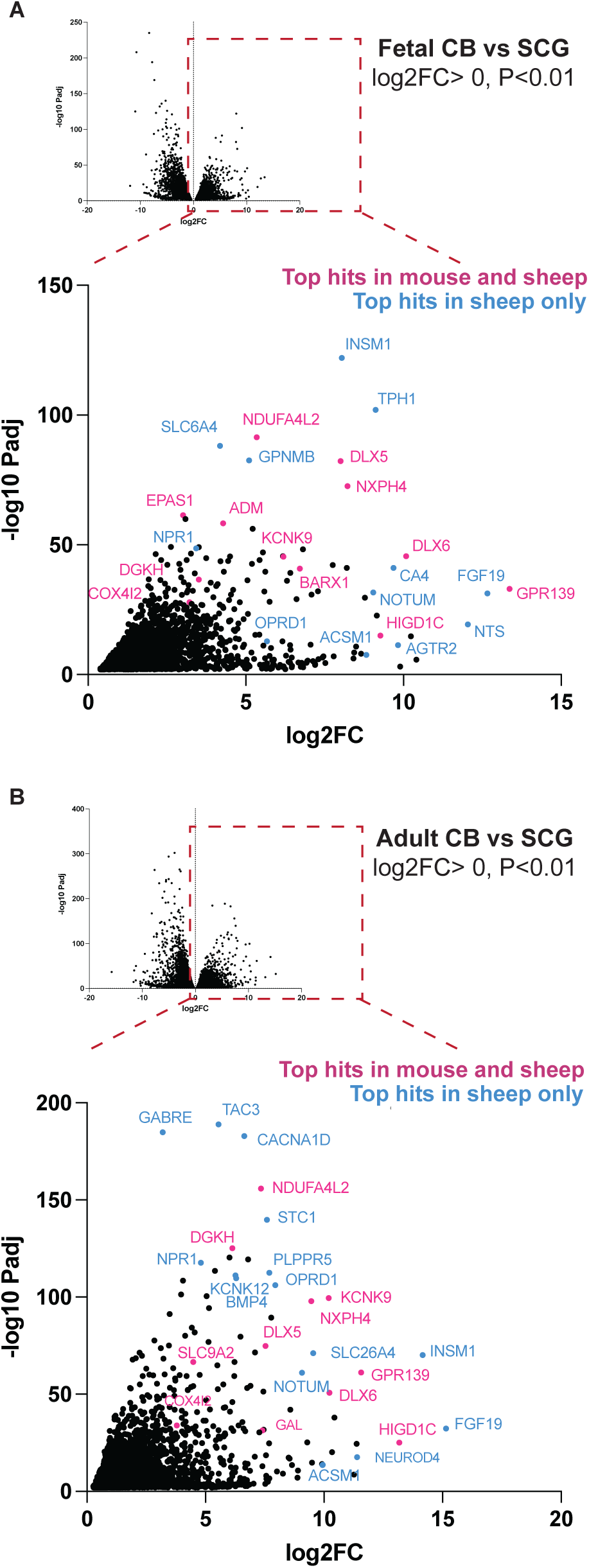
DiDerential gene expression in CB and SCG from the adult sheep and d120 fetus. Volcano plots of the diSerential expression of genes in the CB versus SCG in fetal day 120 (A) and adult sheep (B). In A and B, the lower panel shows the highlighted area (log fold change in expression >0 i.e. up-regulated in CB vs SCG) with annotation of individual genes. Genes highlighted in pink are among the most up-regulated genes in both sheep and mouse CB gene expression datasets, while genes annotated in blue are among the most highly up-regulated genes in the respective sheep CB but are not listed in the top-ranked CB-enriched genes in mouse.

### Features of tissue-specific gene expression identified in the sheep CB

We hypothesized that the greater sequencing depth and power of the sheep dataset, which allows analysis of biological replicates, might reveal additional genes with functionally significant CB-specific expression that were not clearly apparent in previous analyses of pooled mouse tissue. To pursue this, we deepened our examination of genes enriched in the sheep CB by analysis of gene ontology for genes differentially expressed between the CB versus SCG in adult and fetal d120 sheep (Table 2A-D). Considerable similarities between the adult and fetal gene expression were evident between genes up-regulated (table 2A,C) and down-regulated in the CB (table 2B,D). Highly enriched genes up-regulated in both the adult and fetal CB relate to receptor tyrosine kinase/growth factor signalling, along with genes encoding components of the extracellular matrix, which may be related through regulation of growth factor activity. These expression data contrast with features of mouse CB gene expression, where high expression of G protein signalling pathways is observed (Zhou *et al*., 2016; Gao *et al*., 2017), although there are also prominent examples of G protein coupled receptors among the most CB-enriched genes in the sheep (*GPR139, OPRD1, AGTR2, PTGER3*) (Figure 2A-e-g, Figure 3, Table 1C).

**Table 2:**
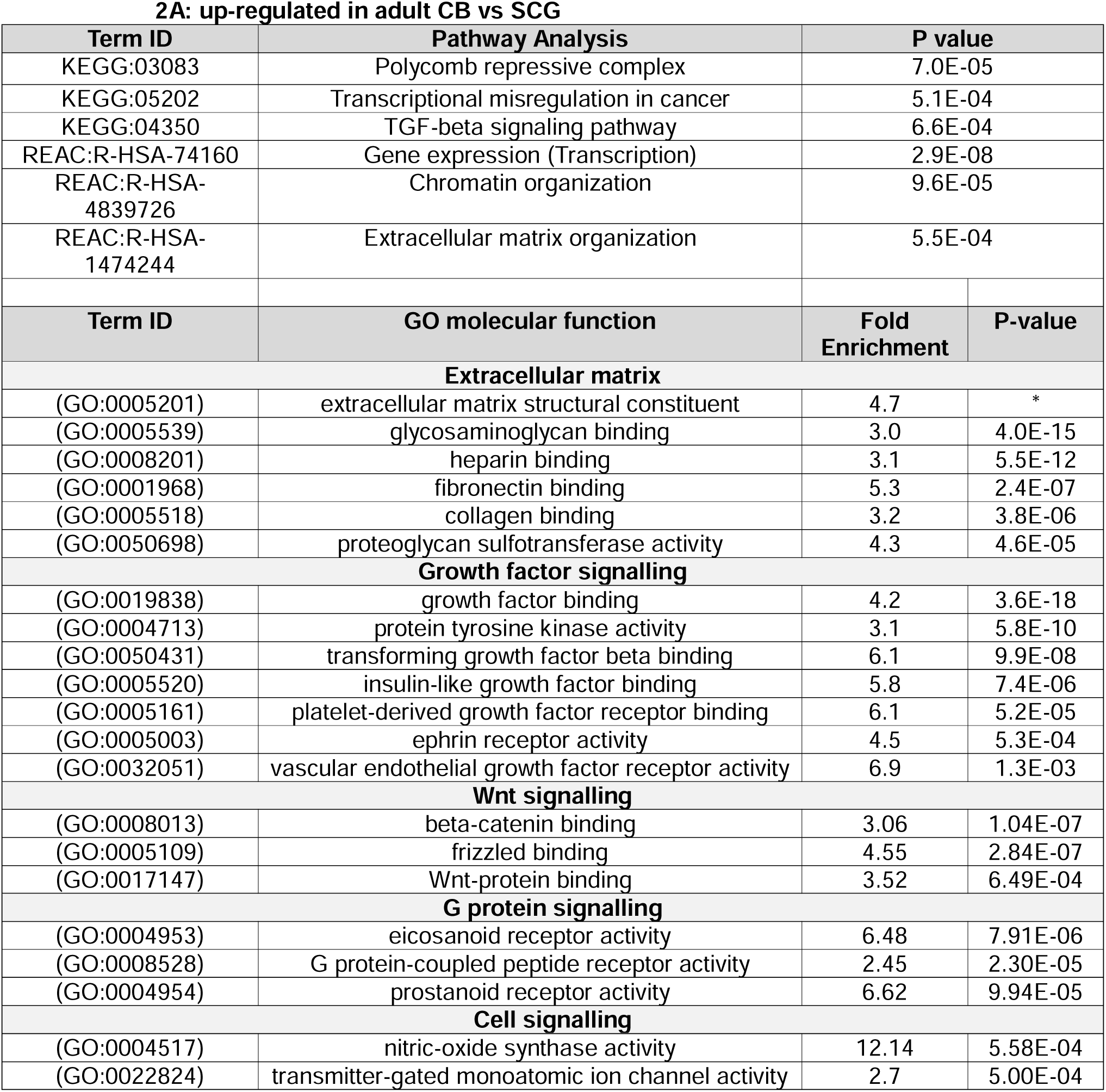
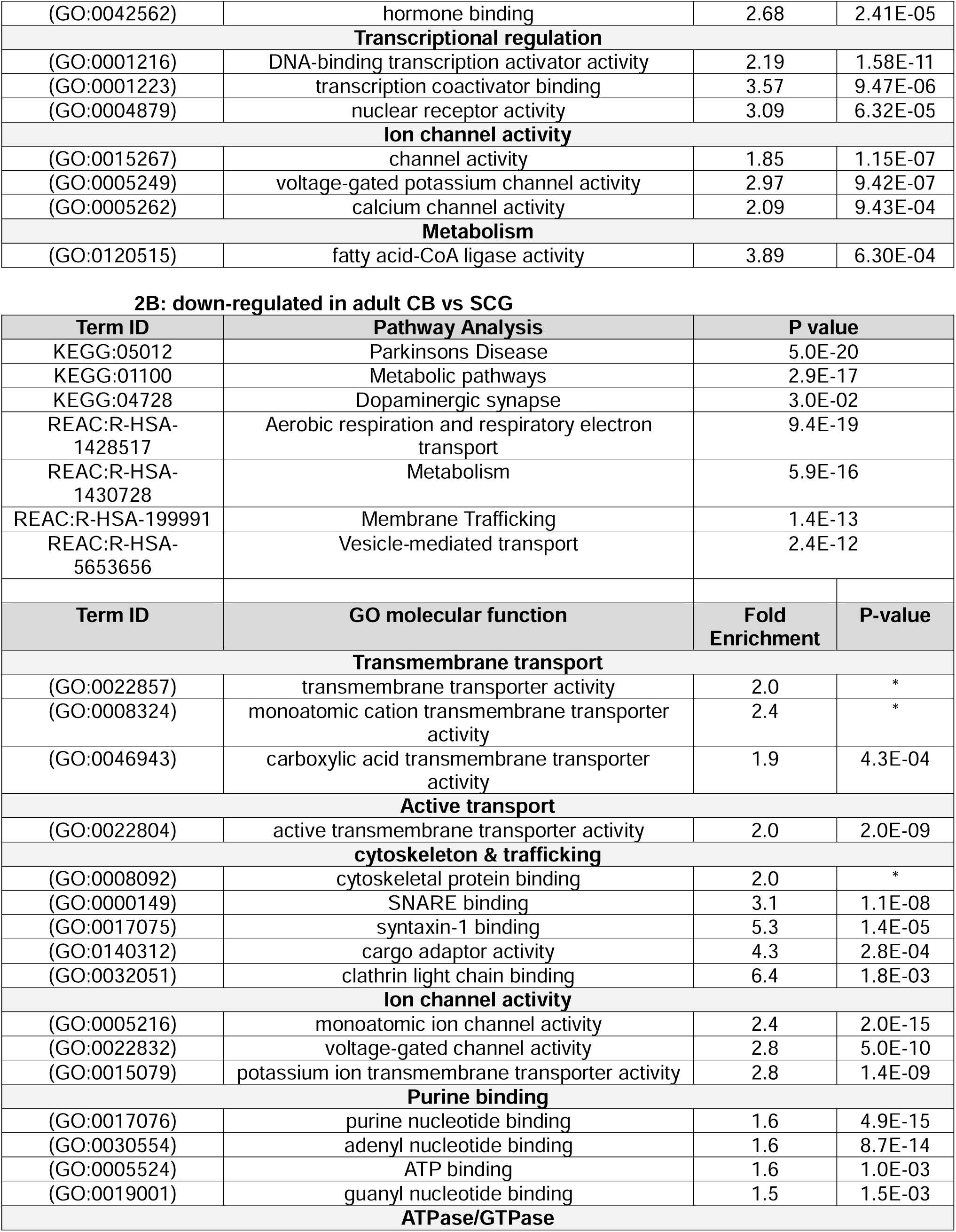
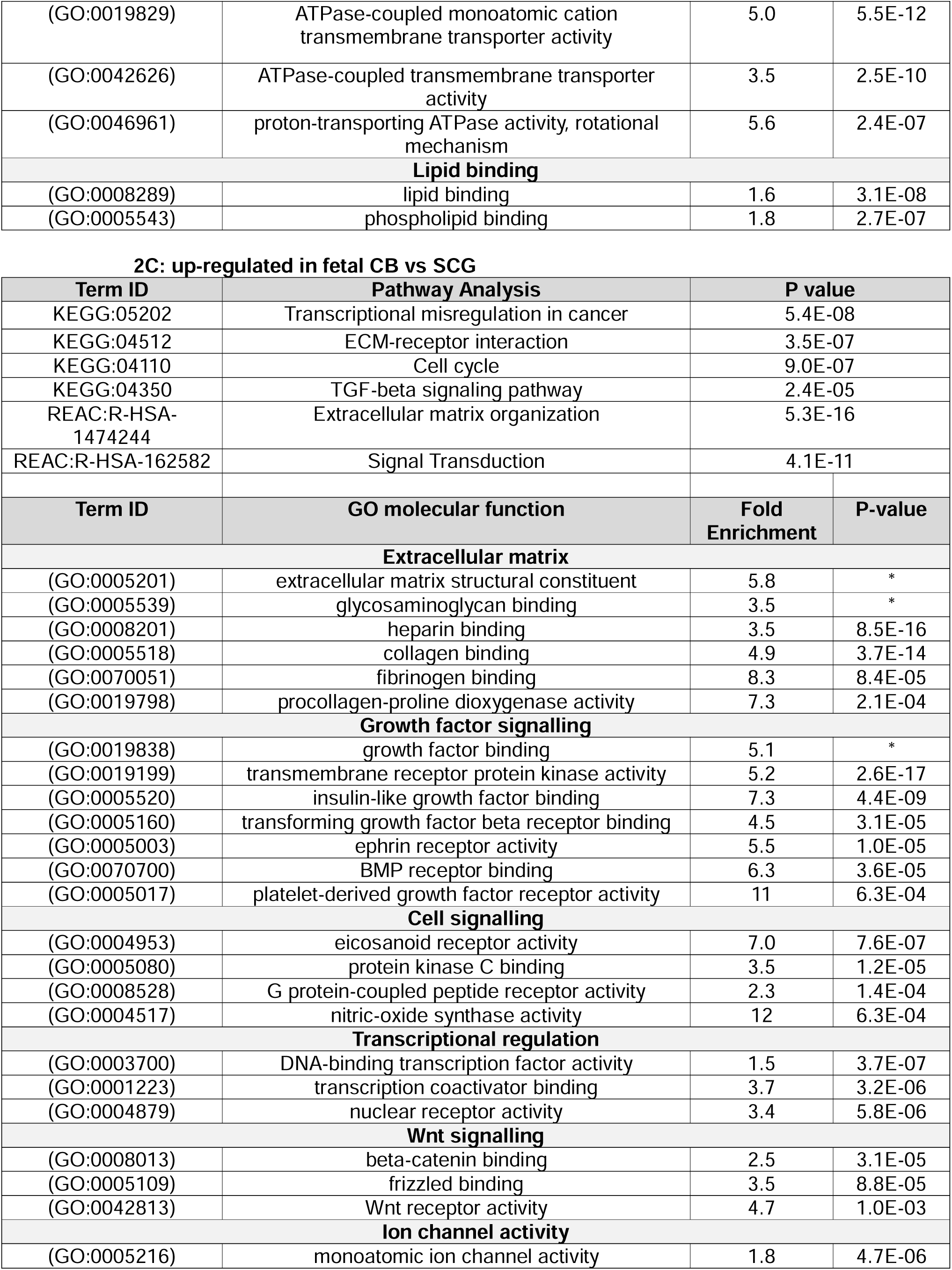
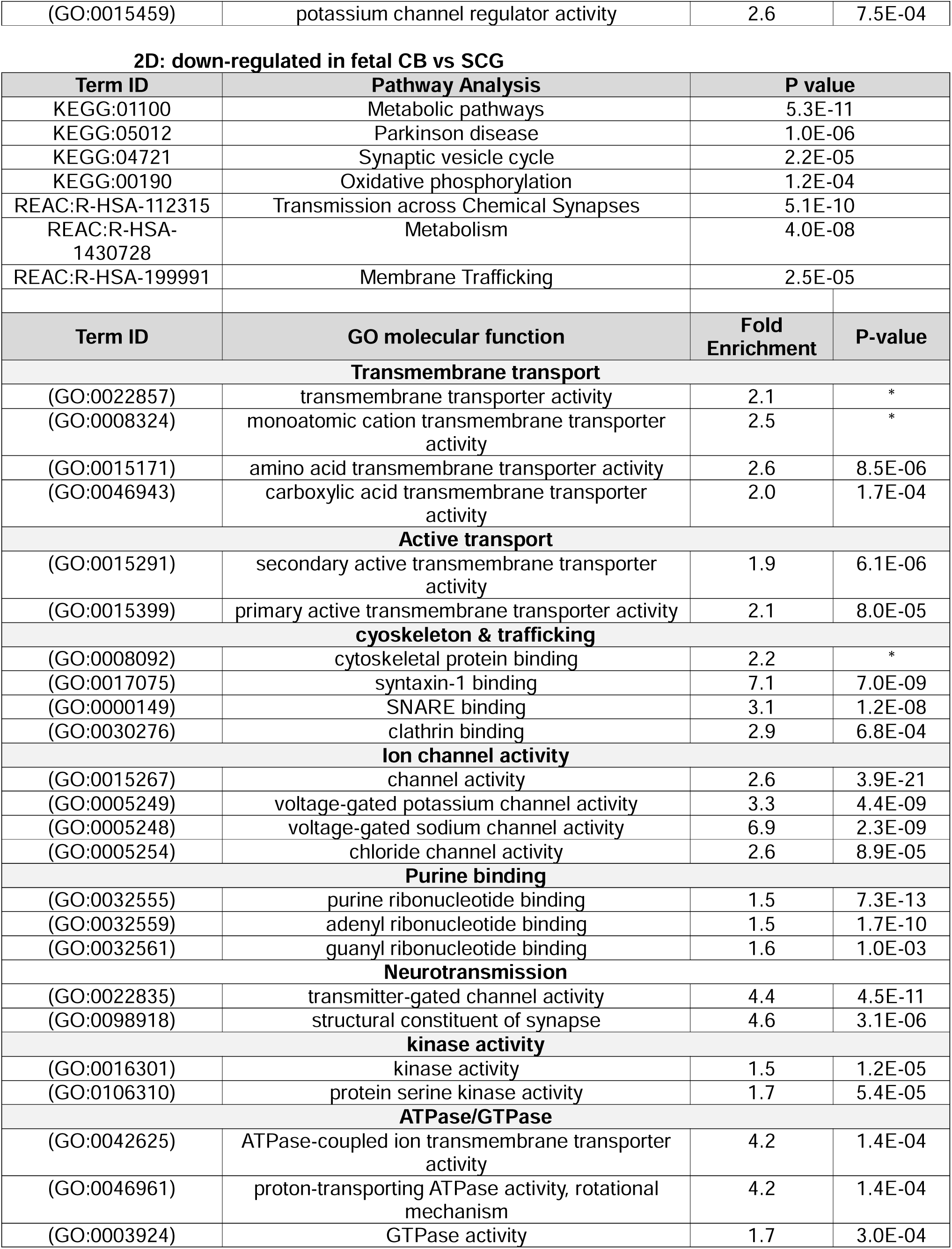
Gene ontology analysis for the most differentially expressed genes in the CB versus SCG of adult and fetal d120 sheep. Tables A-D show the top-ranked biological pathways (KEGG, Reactome) and gene ontology (GO):molecular function terms for the top 2,000 genes showing the greatest up (A) and down-regulation (B) in the CB versus SCG of adult sheep, and genes showing the greatest up (C) and down-regulation (D) in the CB versus SCG of fetal d120 sheep. Terms are shown in order of significance (P value) and organized by category, with similar and redundant terms omitted for clarity. *P value less than 1E-20. All values shown to two significant figures.

Genes enriched in the control tissue, hence relatively down-regulated in the CB, also show features common to adult and fetal CB. Gene ontology analysis of the CB down-regulated genes highlights purine-binding molecules and ATP/GTP-consuming processes such as active transport ATPases, small GTPases, and cellular trafficking (Table 2B,D), suggesting generally low energy turnover. Metabolic pathways are also down-regulated in the CB vs SCG. However, the most CB-enriched genes at each stage include some specific metabolic genes, such as the variant complex IV isoforms (as above), alcohol dehydrogenase (*ADH1C, ADH6*), fatty acid metabolism (*ACSM1*) and other aspects of mitochondrial function (*ATP5MK, APOL3*) (Figure 2A-e-g, Table 1A,C).

Together, the gene expression analysis demonstrates considerable overlap between CB-enriched genes identified in mice and sheep, which is evident even at fetal stages. Analysis of the sheep CB also identifies novel features of CB gene expression, present across developmental stages, including high expression of genes encoding RTK signalling and ECM components. The expression data also provide further evidence of a specific metabolic phenotype in the CB, with down-regulation of genes involved in ATP synthesis and consumption.

### Developmentally regulated genes

We next examined gene expression changes across all time points course studied, during which the CB undergoes significant developmental changes and functional maturation associated with increasing oxygen chemosensitivity. Principal component analysis shows clear separation of the CB and SCG, together with a progressive change in gene expression with time that is evident in both tissues but most marked in the CB (Figure 4A-a).

**Figure 4.**
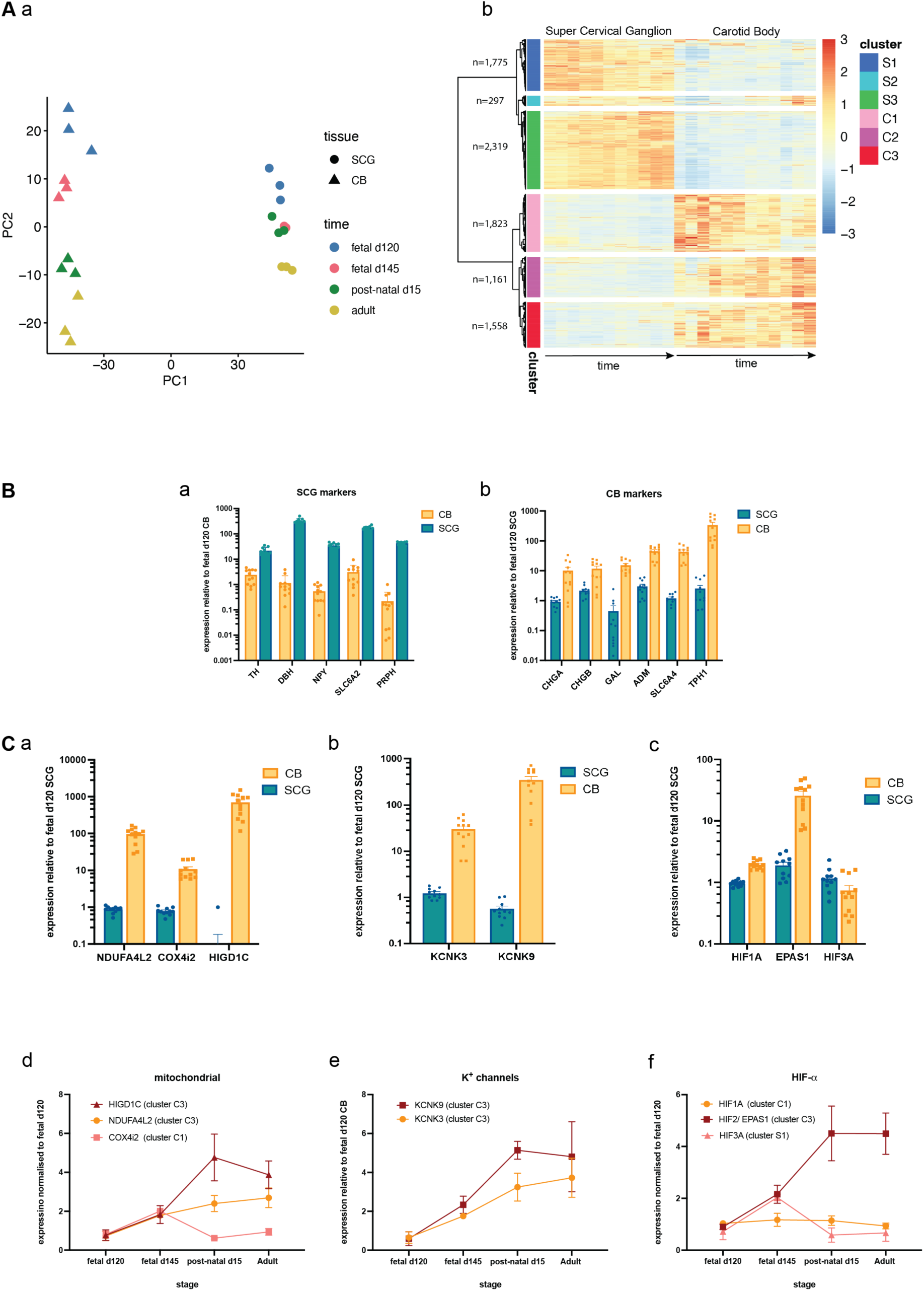
Changes in tissue-specific CB gene expression over developmental time. (A-a) Principal component analysis of gene expression profiles of each sample, including CB (triangle) and SCG (circles) from fetal day 120 (blue), fetal day 145 (pink), post-natal day 15 (green), and adult (yellow). (A-b) Heat map illustrating the hierarchical clustering analysis on all genes showing significant diSerential expression at any time point (n=8,933). The graded colour scale indicates the z score for relative expression of a given gene scaled across each tissue and developmental time point. (B) Expression of tissue specific markers (indicated by gene names) for SCG (blue) and CB (orange), normalized to expression levels in the fetal day 120 CB (a) or SCG (b). Note that, as expression in the reference tissue changes over time, the mean may not equal to 1. (C) Expression of genes implicated in oxygen chemosensing (a-subunits and regulators of complex IV; b-TASK1 (KCNK3) and TASK3 (KCNK9) potassium channels; c-HIF-⍺ isoforms) in CB and SCG samples, normalized to the fetal day 120 SCG. (d-f) Mean expression of these genes in CB samples at each time point, normalized to fetal d120 CB. Bar charts show mean expression (+/-SEM) with dots showing values for individual samples. Graphs depicting time courses show mean expression of the specified gene (+/-SEM) at each time point.

This monotonic change in overall gene expression profile most likely reflects multiple underlying processes associated with potentially complex gene expression patterns that increase or decrease over time. To study this in more detail we performed a hierarchical clustering analysis across all time points, incorporating genes that showed significant differential gene expression between tissues at any time point (see *Methods*). This yielded 6 clusters: three comprising genes that are CB-enriched (C1, C2, C3) and three with SCG-enriched genes (S1, S2, S3). Cluster C1 and S1 genes are predominantly enriched at early/fetal time points, while C3 and S3 genes increase with development and maturation. Genes in clusters C2 and S2 show more complex, intermediate temporal patterns of expression (Figure 4A-b).

To evaluate the results of the clustering analysis, we fist examined the cluster allocation of known marker genes for each tissue. Neuronal markers (*PRPH*) and genes related to sympathetic neurotransmitters (noradrenaline and neuropeptide Y) mapped to SCG-enriched cluster, specifically cluster S3 (Figure 4B-a). Tyrosine hydroxylase (*TH*), catalysing dopamine biosynthesis, is highly expressed in both tissues (as also seen at the protein level (Figure 1C)) but correlates more with sympathetic development (cluster S3). Other CB markers, including chromogranin secretory proteins, signalling neuropeptides, and genes related to serotonin production, were all highly up-regulated in the sheep CB and map to CB-enriched clusters C1 (*GAL, TPH1*), C2 (*ADM*), or C3 (*CHGA, CHGB, SLC6A4*) (Figure 4B-b).

We next examined temporal changes in expression of genes already implicated in the oxygen chemosensitivity of chemosensory cells (*KCNK3/9, COX4I2, NDUFA4L2, HIGD1C, EPAS1/HIF2*). In addition to the high CB-specific expression of these genes (Figure 2A-d, Figure 3B, 4C-a-c), we also see striking ontogenic changes in expression. Most of the selected “chemosensitivity genes” show progressive CB-enrichment, mapping to cluster C3. This includes TASK1/3 potassium channels (*KCNK3/9*, Figure 4C-e) and variant isoforms of proteins contributing to complex IV function (*NDUF4AL2, HIGD1C*), except for *COX4I2*, which peaks in fetal stages (cluster C1) so may be important during pre-natal development (Figure 4 C-d). *COX8B*, which is up-regulated in the mouse CB, does not show significant differential expression in the sheep CB (data not shown). The HIF pathway has significant roles in regulating CB development and function and, as in mice, we find specific enrichment of the HIF-2α isoform (*EPAS1*) in sheep CB, which increases with development (cluster C3, Figure 4C-f).

Together, the clustering analysis supports the hypothesis that the expression of genes mediating the oxygen chemosensitivity response correlates with the time course of the physiological maturation of this response and are enriched in the CB-maturation cluster (C3). Hence, we wished to examine this cluster of genes more closely. There are some individual genes in C3 that are interesting examples due to potential clinical links with pharmacological and pathological variation in CB function. The delta opioid receptor (*OPRD1*) is one of the most up-regulated genes in the adult CB (Figure 2A-e, Figure 3B) and shows developmentally increasing expression, in contrast to other opioid receptors which are not CB-enriched (Figure 5A). Opioid receptor agonists cause respiratory depression, in part through suppression of CB chemoreceptor-mediated hypoxic ventilatory response. These data indicate that this is likely to be predominantly through the delta receptor isoform, consistent with findings from pharmacological studies in cats (Kirby & McQueen, 1986) . Natriuretic peptide receptor 1 (*NPR1*) is one of the most CB-enriched genes in adult and fetal CB (Figure 3B), with a developmentally increasing expression pattern (C3), in contrast to related isoforms of this receptor (Figure 5B). This is interesting given evidence from both clinical and pre-clinical studies for CB hyperactivity and associated sympathetic activation in heart failure (Schultz *et al*., 2013), a condition in which circulating natriuretic peptides are pathologically elevated. The mechanism of CB sensitisation in heart failure is unknown, but these gene expression data show the potential for direct action of natriuretic peptides on CB cells, which may have a physiological function that is dysregulated by pathological excess of natriuretic peptides.

**Figure 5.**
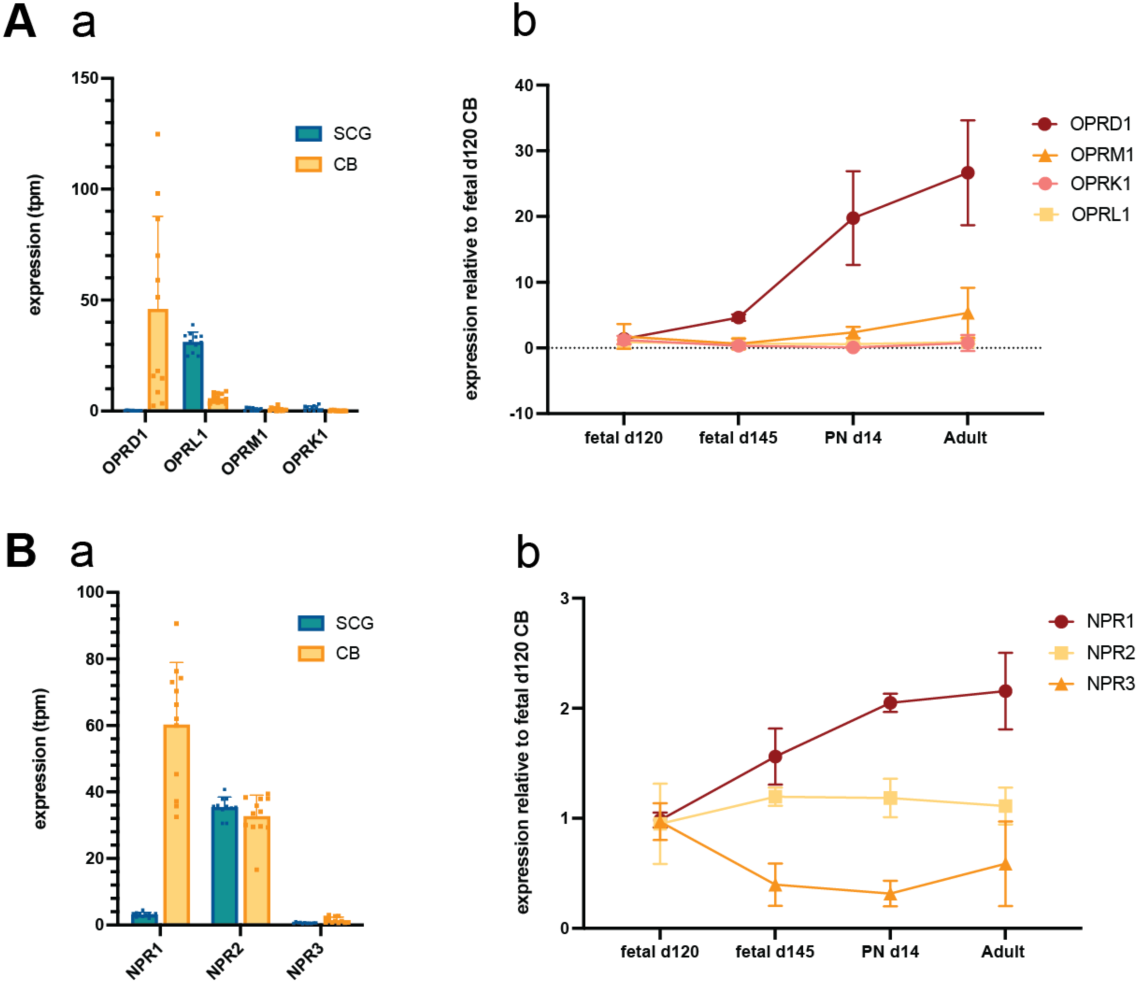
Genes encoding opioid and natriuretic peptide receptors exhibit CB-enriched expression that correlates with CB maturation. Expression of opioid receptors (A) and natriuretic peptide receptors (B), showing absolute expression levels (tpm) in SCG (blue) and CB (yellow) samples (A-a, B-a) and mean expression in CB samples at each time point (A-b, B-b), normalized to fetal d120 CB. Bar charts show mean expression (+/-SEM) with dots showing values for individual samples. Graphs depicting time courses show mean expression of the specified gene (+/-SEM) at each time point.

### G proteins and Diacylglycerol metabolism

Gene ontology analysis of C3 genes shows enrichment of glycosaminoglycan metabolism, potassium channels, and diacylglycerol (DAG) and IP3 signalling (Figure 6). In the mouse CB, DAG signalling was found to be enriched in the context of genes encoding Gq proteins and Gq-coupled receptors (Zhou *et al*., 2016; Gao *et al*., 2017). However, we do not find CB-enrichment of the Gq class of G proteins (*GNAQ, GNA11*), nor several other G protein signalling genes that are strikingly highly expressed in mouse CB (*GNAS, RGS4*) (Figure 7A-a). An exception to this is *RGS5*, which is one of the most highly expressed genes in the C3 cluster (Table 3), showing marked up-regulation in the CB that is progressive with time (Figure 7A-a-b). Of the primary phospholipase C enzymes (generating DAG and IP3 from membrane phospholipids), only the gamma isoform (*PLCG1*), mediating receptor tyrosine kinase signalling, is CB-enriched, mapping to cluster C3, while the isoforms linked to G protein coupled receptors (GPCR) do not show significant differential expression (Figure 7A-c). The most striking DAG/IP3 related gene in C3 is diacylglycerol kinase eta (*DGKH*), which is one of the most highly expressed genes in this cluster (Table 3). *Dgkh* is also highly expressed in the mouse CB, together with *Dgkk* and *Dgkg*, but *DGKH* is the only diacylglycerol kinase significantly enriched in the sheep CB (Figure 6B-a). We have confirmed this high expression at the protein level (Figure 7B-c), along with a clear pattern of developmentally progressive transcript expression (Figure 7B-b). Interestingly, *PLPPR3* and *PLPPR5*, encoding phosphatidic acid phosphatases (PAP) that generate DAG, are also in cluster C3. Analysing the clusters of other PAP genes identifies four in CB-enriched clusters: *PLPP3, PLPPR1, PLPPR3* and *PLPPR5*. However, only *PLPPR5* shows a clear pattern of developmentally increased expression (Figure 7C-a-b), while *PLPPR1* and *PLPP3* cluster in C1, and *PLPPR3* only increases relative to decreasing SCG-expression (data not shown).

**Table 3:** Cluster C3 genes with the highest mean CB expression. The top 20 genes in the CB maturation cluster (C3), ranked by absolute expression (tpm) averaged across CB samples from all stages (fetal d120, fetal d145, post-natal d15, and adult).

| Gene name | Description | average CB expression (tpm) |
| --- | --- | --- |
| *CHGB* | Chromogranin B/ Secretogranin B | 4168 |
| *RGS5* | Regulator of G protein signalling 5 | 2527 |
| *EPAS1* | Endothelial PAS Domain Protein 1/ HIF-2. | 1327 |
| *H3F3A* | H3.3 Histone A | 1088 |
| *NDUFA4L2* | NDUFA4 Mitochondrial Complex Associated Like 2 | 880 |
| *CHGA* | Chromogranin A | 825 |
| *LOC114112700* | Translation initiation factor IF-2-like. | 788 |
| *HSPA1A* | Heat Shock Protein Family A (Hsp70) Member 1A | 642 |
| *DGKH* | Diacylglycerol Kinase Eta | 625 |
| *TAC3* | Tachykinin Precursor 3 | 603 |
| *HIGD1C* | HIG1 Hypoxia Inducible Domain Family Member 1C | 547 |
| *SCG2* | Secretogranin II | 503 |
| *BEX2* | BEX2, Brain Expressed X-Linked 2 | 478 |
| *VEGFA* | Vascular Endothelial Growth Factor A | 377 |
| *FAM162B* | Family With Sequence Similarity 162 Member B | 318 |
| *ADCYAP1R1* | Adenylate Cyclase Activating Polypeptide 1 (Pituitary) Receptor Type I | 308 |
| *INSM1* | Insulinoma-Associated Protein 1 | 298 |
| *CA4* | Carbonic Anhydrase 4 | 276 |
| *HSPA1B* | Heat shock 70 kDa protein 1B | 266 |
| *TMT1A* | Thiol Methyltransferase 1A | 249 |

**Figure 6.**
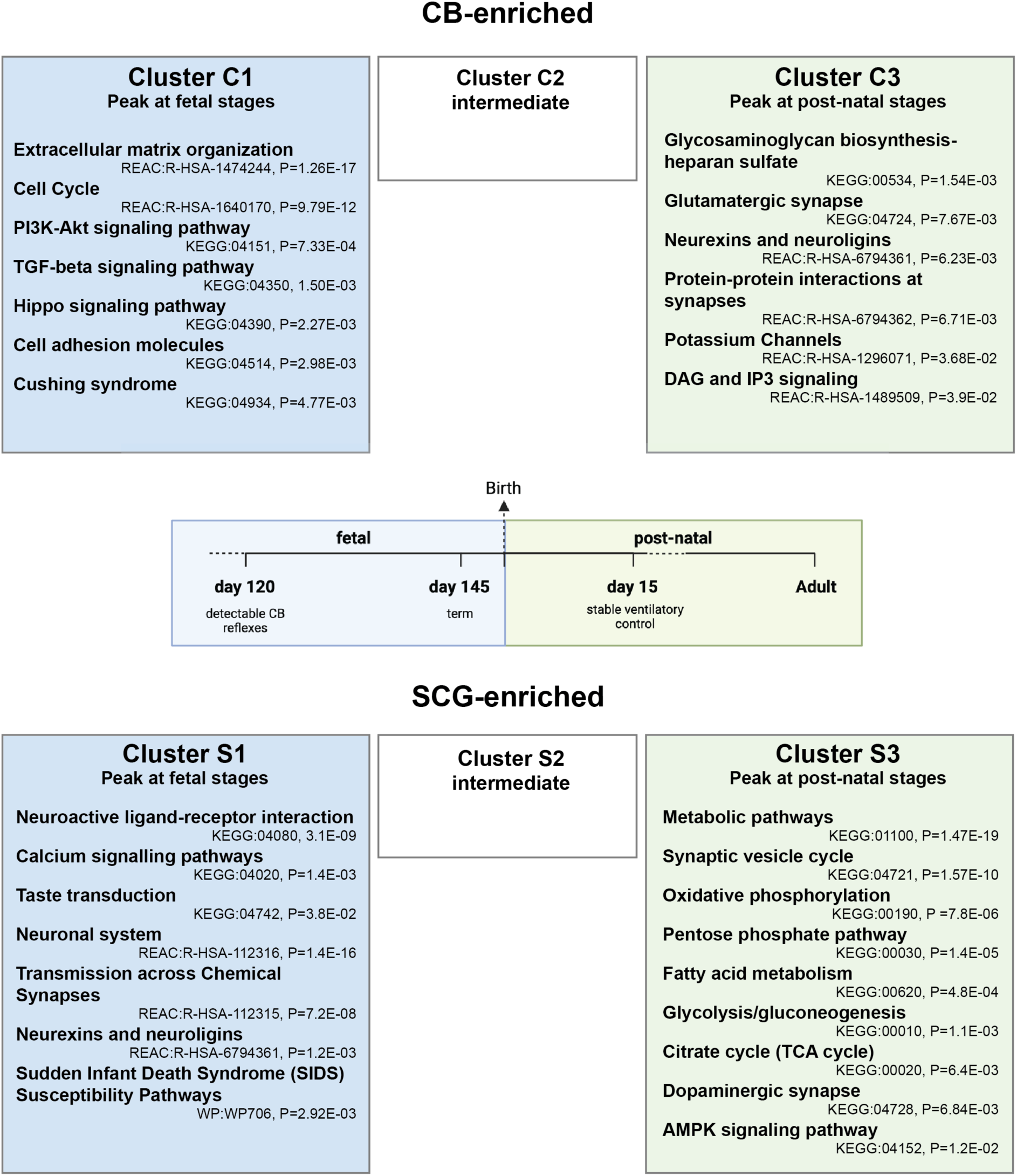
Biological pathways enriched in tissue-specific developmental clusters of CB gene expression. The most significantly enriched biological pathways in each of the four main gene expression clusters that are predominantly up-regulated in the CB (C1, C3) or SCG (S1, S3) at fetal (C1, S1), or post-natal stages (C3, S3). Pathway analysis by KEGG, Reactome (REAC), or Wikipathways (WP), with the listed pathways ordered by significance and selected to reduce redundant or related terms.

**Figure 7.**
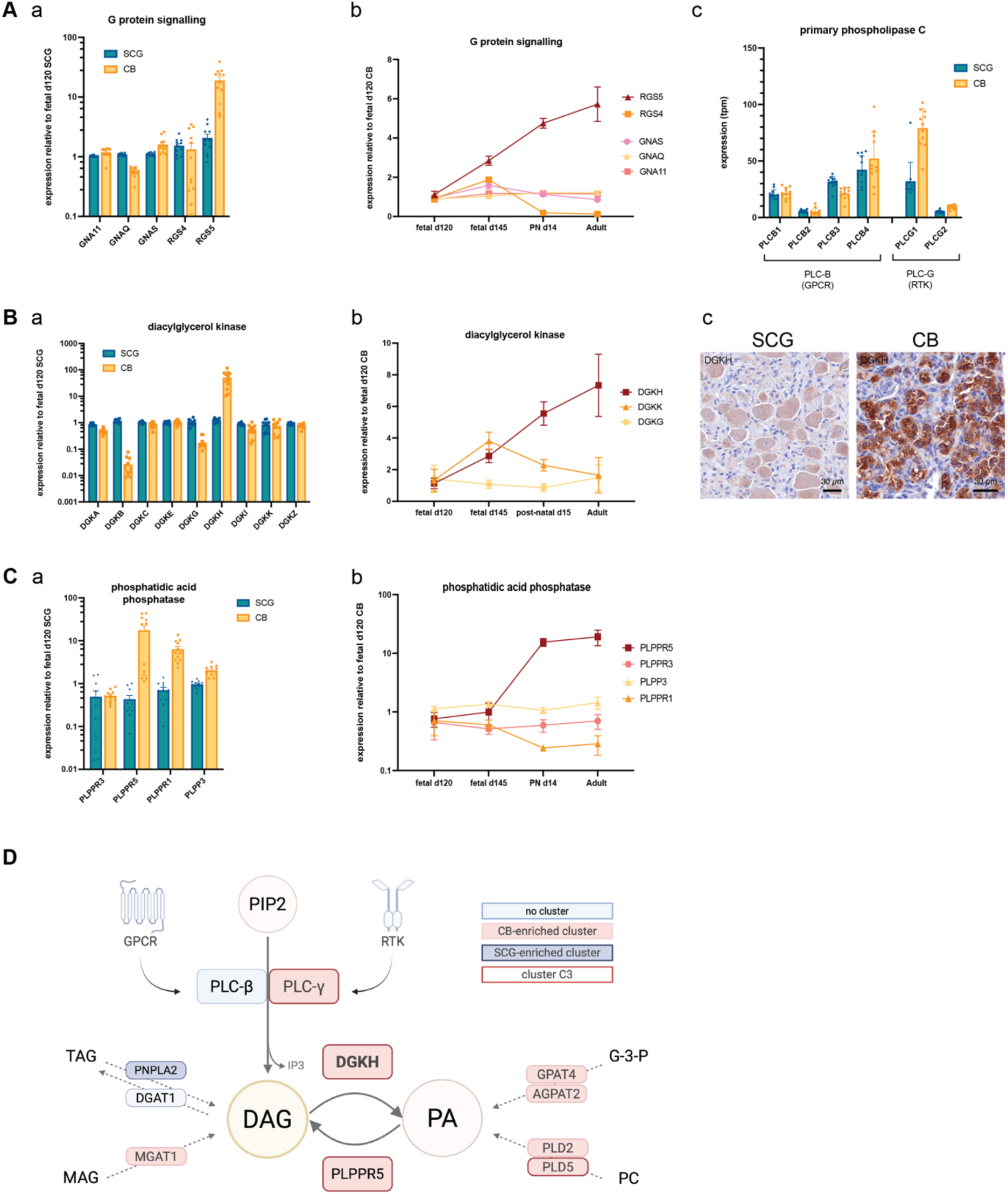
Expression of genes regulating diacylglycerol signalling during CB maturation. (A)(a-b) Relative expression of selected G proteins and regulators of G protein signalling: (a) mean expression in SCG (blue) and CB (yellow) samples normalized to fetal d120 SCG; (b) mean expression in CB samples at each time point normalized to CB fetal d120. (c) Relative expression of phospholipase C isoforms, showing mean expression each gene in SCG (blue) and CB (yellow) samples normalized to fetal d120 SCG. (B) Relative expression of diacylglycerol kinase isoforms, showing (a) mean expression in SCG (blue) and CB (yellow) samples normalized to fetal d120 SCG and (b) mean expression in CB samples at each time point normalized to fetal d120 CB. (c) Immunostaining for DGKH in adult SCG and CB (scale bar= 30 um). (C) Relative expression of phosphatidic acid phosphatase isoforms, showing (a) mean expression in SCG (blue) and CB (yellow) samples normalized to fetal d120 SCG and (b) mean expression in CB samples at each time point normalized to fetal d120 CB. Bar charts show mean expression (+/-SEM) with dots indicating values for individual samples. Graphs depicting time courses show mean expression of the specified gene (+/-SEM) at each time point. (D) Schematic diagram of diacylglyerol metabolism including: membrane phospholipid metabolism regulated by G protein couple receptors (GPCR) or receptor tyrosine kinases (RTK); fatty acid metabolism via triacylglycerol (TAG) or monoacylglycerol (MAG); the glycerol-3-phosphate (G-3-P) pathway; and phosphatidyl choline (PC) metabolism. Enzymes/receptors in each pathway are indicated by the gene names in boxes highlighted by tissue/time gene expression cluster (pink=CB-enriched, red border = cluster C3, dark blue=SCG-enriched, light blue=no cluster). Diagram created using Biorender.

Multiple metabolic pathways can generate phosphatidic acid and diacylglycerol, including the glycerol-3-phosphate pathway, fatty acid, and phosphatidyl choline metabolism. Genes encoding metabolic enzymes in each of these pathways map to CB-enriched expression clusters (Figure 7D), suggesting multiple contributions to the DAG pool that may vary by developmental stage.

Taken together, this suggests that a key, conserved feature of mature CB gene expression centres on the regulation of diacylglycerol, with tissue-specific, developmentally increased expression of genes catalysing the generation of diacylglycerol (*PLCG1, PLPPR5*) and its conversion to phosphatidic acid (*DGKH*). The nature of upstream regulators/ receptors that feed into this pathway may vary between species, with GPCR being dominant in mouse, while receptor tyrosine kinases are more highly expressed in the sheep CB.

### Energy metabolism

While our primary focus is on genes showing CB-enriched expression, genes that that are down-regulated in a tissue specific manner may also be important for generating cellular phenotypes. Strikingly, genes regulating many aspects of bioenergetic metabolism, including the core metabolic pathways of glycolysis, the tricarboxylic acid (TCA) cycle, and oxidative phosphorylation, are consistently down-regulated in CB samples but are progressively enriched in the SCG with maturation (cluster S3, Figure 6), with the differential expression of these genes being most marked in the adult (Table 3B). Gene set enrichment analysis confirms these effects (Figure 8A), also showing down-regulation of the pentose phosphate pathway and fatty acid metabolism in the CB, but relative sparing of amino acid metabolism. The magnitude of these effects at the level of individual genes is illustrated by volcano plots of gene sets for glycolysis, TCA cycle, and oxidative phosphorylation (Figure 8B), which also highlight a small number of genes that defy this trend. As previously noted, these include the variant isoforms of complex IV (*NDUFA4AL2, COX4I2*), and pyruvate carboxylase (*PC*), which is also up-regulated in the mouse CB, possibly mediating anaplerotic fuelling of the TCA cycle (Gao *et al*., 2017). In addition, we found CB-enrichment of *PCK1* (phosphoenolpyruvate carboxykinase 1), which may also contribute to anaplerosis, and SUCLG2, the isoform of Succinate-CoA Ligase that specifically generates GTP. Expression of the ATP-specific *SUCLA2* is suppressed in the CB (data not shown). However, the broader picture suggests that both forms of molecular energy currency (ATP/ GTP) are likely to be produced at lower levels in the context of widespread metabolic suppression in the CB. Interestingly, this occurs in parallel with down-regulation of ATP/GTP binding proteins and energy consuming processes, as highlighted in the differential expression analysis of the adult and fetal CB vs SCG (Table 3B,D). Together, this builds a picture of a bioenergetic phenotype that extends beyond the previously established specific variations in CB metabolism, such as atypical complex IV.

**Figure 8.**
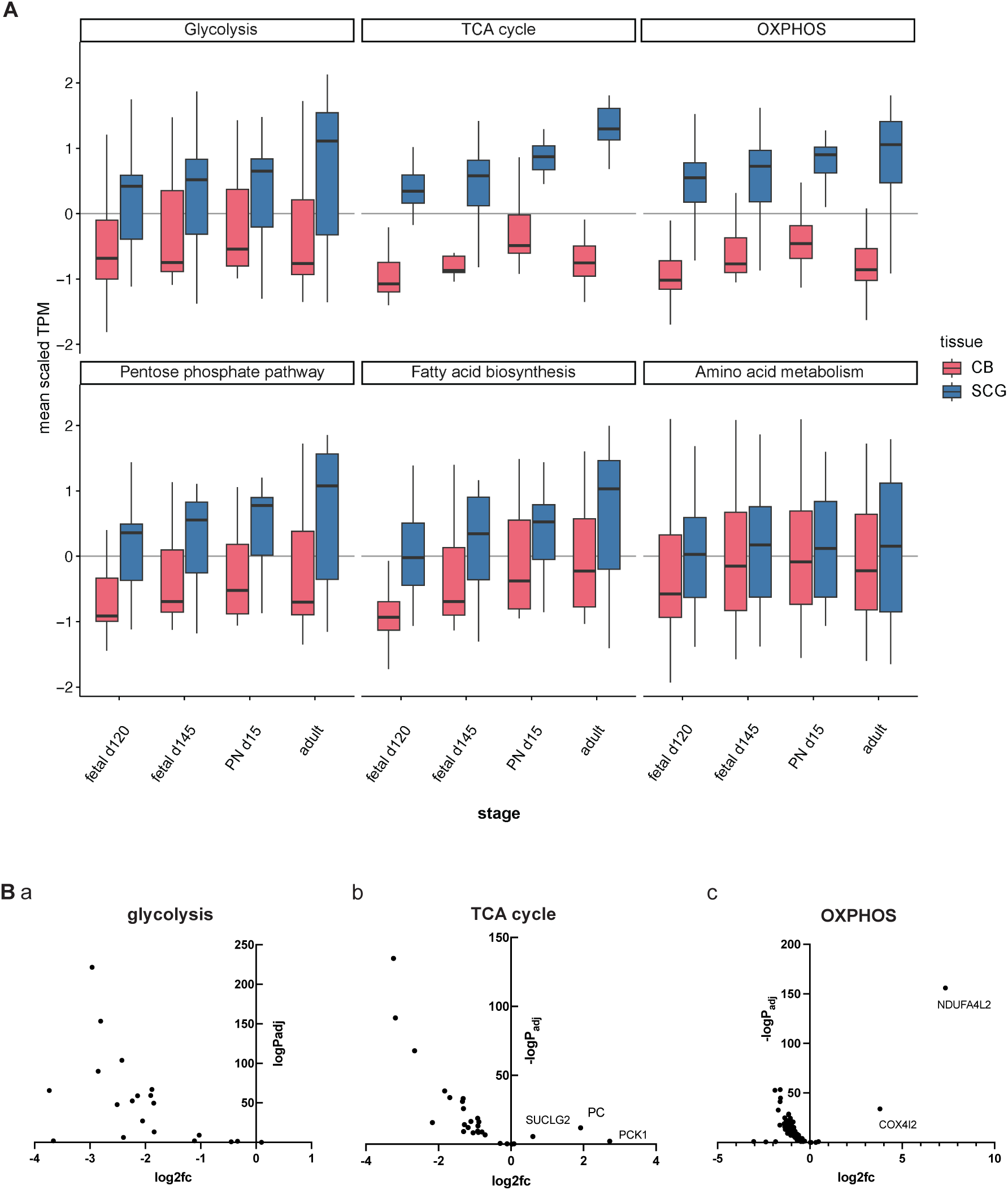
Down-regulation of genes encoding bioenergetic pathways in the CB versus SCG. (A) Gene set enrichment analysis for metabolic pathways in CB (red) vs SCG (blue) at each time point, expression z score for each developmental stage (+/-interquartile range (boxes) and full range). (B) Volcano plots of diSerential expression in adult CB versus SCG for glycolysis (a), tricarboxylic acid cycle (b) and oxidative phosphorylation genes (c). The annotated genes are upregulated in CB versus SCG.

### Extracellular matrix

Genes encoding extracellular matrix (ECM) components are consistently a dominant feature of CB gene expression in the sheep, being amongst the most highly enriched genes in both the adult and day 120 fetal CB (Table 3A,C). Of genes encoding structural components of the ECM, including the fibrous proteins collagen, elastin, and fibronectin, and the proteoglycans (total n=87), 40 genes are in cluster C1, and, overall 58/87 ECM genes map to a CB-enriched cluster (C1, C2, C3 – Figure 9A). The cluster analysis indicates that ECM genes predominantly exhibit a fetal-enriched expression pattern (C1), as illustrated by genes encoding type I collagen and elastin (Figure 9B-a-b). The pattern is observed widely across ECM genes, but most striking for genes encoding collagens, with 23 of 41 annotated collagen genes found in cluster C1. At the protein level, Masson Trichrome staining confirms a high density of collagen surrounding the nested glomeruli of CB type I cells, in contrast to the far less prominent ECM background of the SCG (Figure 9C). This dense CB ECM does not appear significantly different in the adult versus the fetal CB, so may be established early with relatively little turnover. Interestingly, enzymes regulating biosynthesis and metabolism of heparan sulphate/ glycosaminoglycans (not included in Figure 8A) are significantly enriched in cluster C3 (Figure 5). These exhibit a developmentally increasing expression pattern, as illustrated by *B3GAT2* and *HS6ST2* (Figure 9B-c-d), so the ECM composition may evolve with CB maturation.

**Figure 9.**
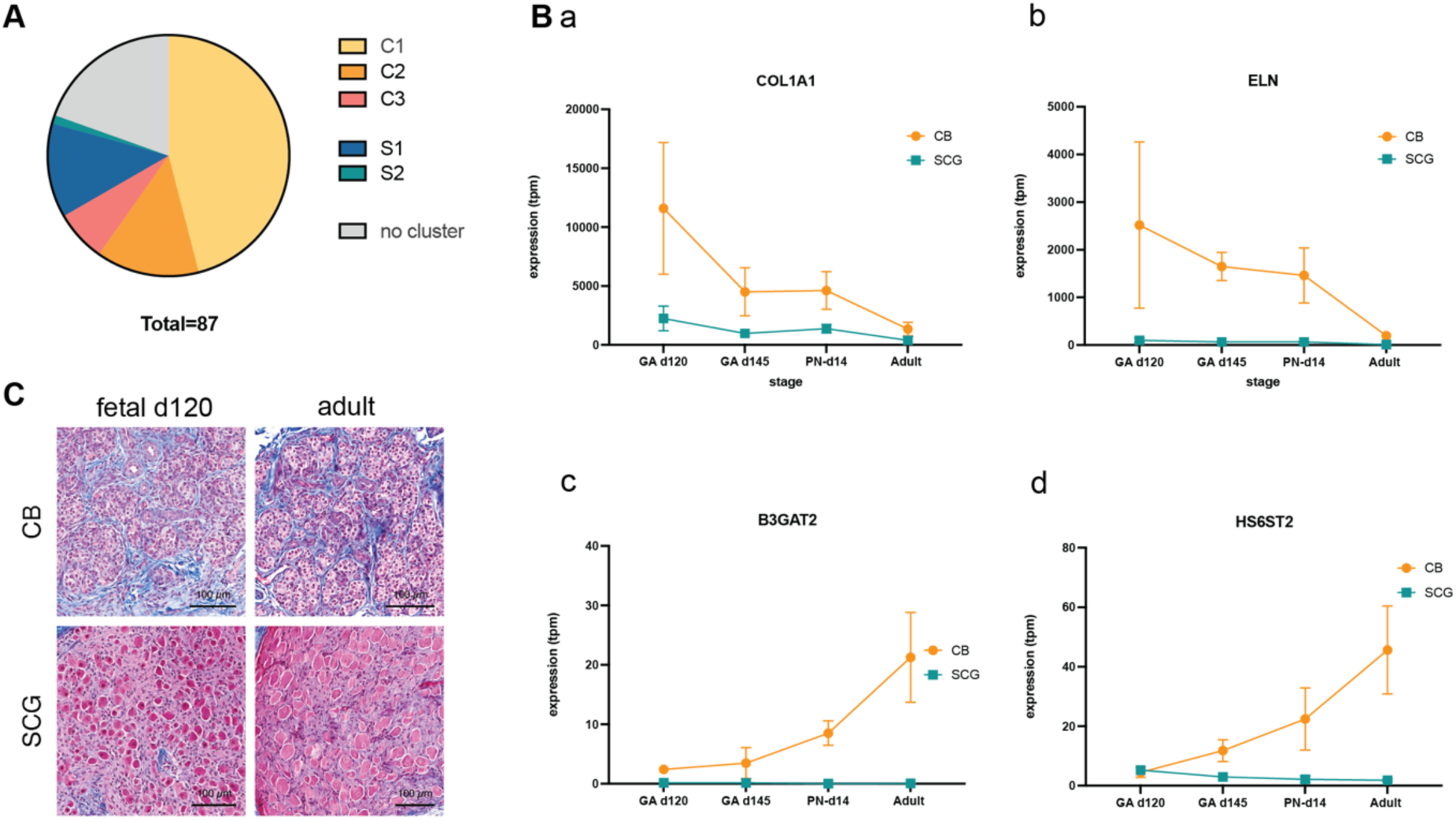
Extracellular matrix gene expression in the CB versus SCG. (A) Developmental gene expression cluster of extracellular matrix components (n=87 genes), including collagens, elastin, laminins, tenascin, fibronectin, and the proteoglycans. Genes that did not meet the criteria for cluster analysis are shown in grey (no cluster). Note, most of 87 genes map to cluster C1, but none map to cluster S3, hence this is not shown. (B) Expression of extracellular matrix genes (collagen A1 (a) and elastin (b)) and enzymes for glycosaminoglycan biosynthesis (Beta-1,3-Glucuronyltransferase 2 (c) and Heparan Sulfate 6-O-Sulfotransferase 2 (d) showing mean expression in CB (orange) or SCG samples (blue) at each time point (mean expression (tpm) +/-SEM). (C) Masson Trichrome staining of CB and SCG sections from fetal day 120 and adult sheep. Collagen is stained blue, scale bar is 100 uM.

### Transcription factors

Having identified clusters of developmentally regulated gene expression patterns we were interested in which transcription factors might mediate these programs, with associated roles in CB development and maturation of chemosensitivity. We identified the transcription factors in each of the tissue-time gene expression clusters and examined the temporal expression pattern for individual examples (Figure 10A).

**Figure 10.**
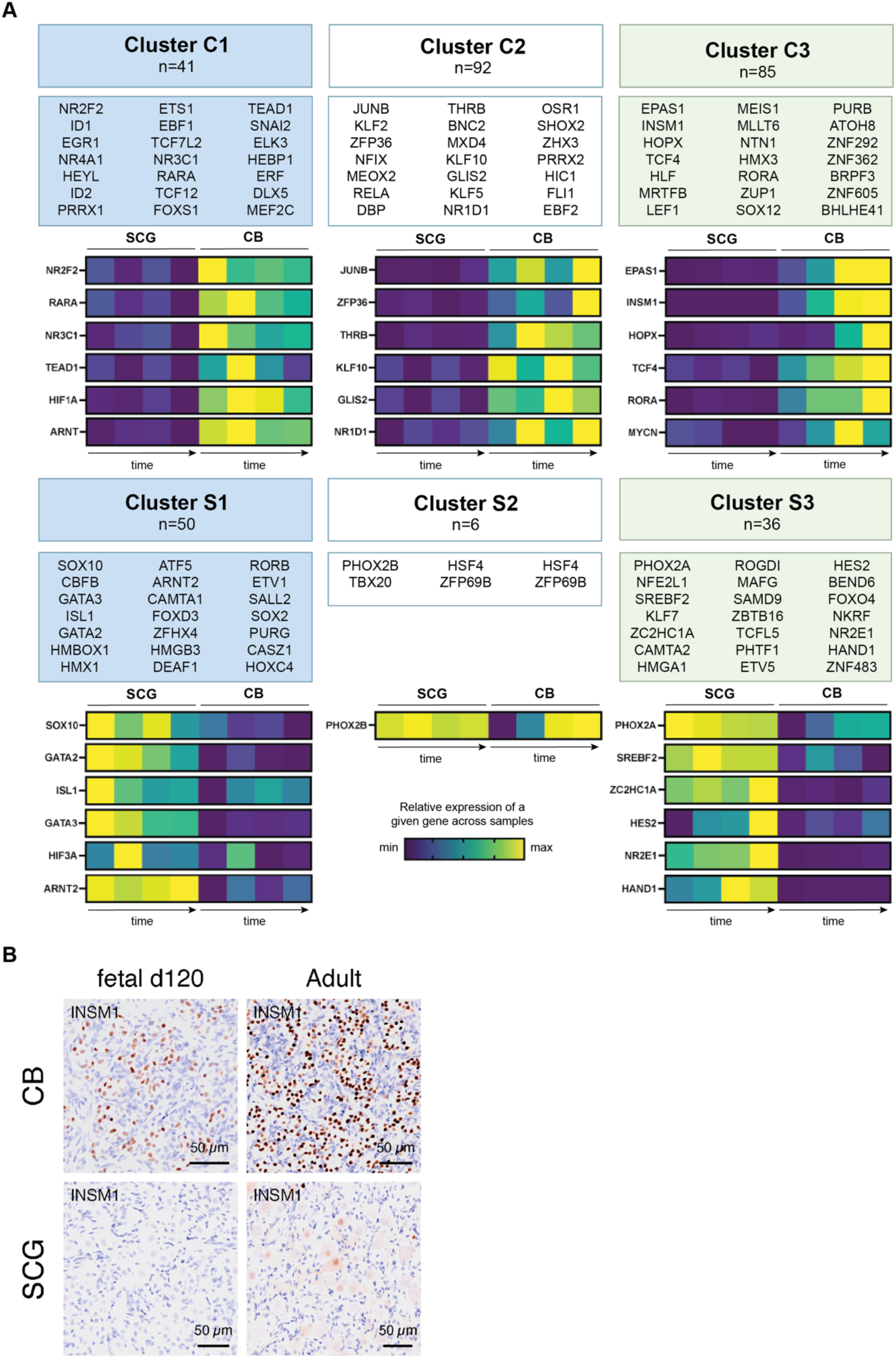
Transcription factors with distinct temporal and tissue-specific expression patterns. (A) Transcription factors in each gene expression cluster. For each cluster, the top 21 most highly expressed (where n>21) are listed in order of average expression in the fetal d120 CB (C1), adult CB (C3), all CB samples (C2), fetal d120 SCG (S1), adult SCG (S3) or all SCG samples (S2) (in each case aiming to list in order of maximal expression). Also shown are individual heat maps for a selection of transcription factors illustrating interesting patterns of developmental expression. The graded colour scale indicates relative expression of a given gene scaled across each tissue and developmental time point. Each square represents mean expression in that tissue and time point. The arrows marked “time” indicate the progression from fetal d120, to fetal d145, post-natal d15, and adult. (B) Immunostaining for INSM1 in CB and SCG from fetal d120 and adult sheep. Scale bar = 50 um

Overall, transcription factors are more frequent in CB-enriched clusters (Figure 10A), many with complex temporal expression patterns. The HIF transcription factors have interesting developmental expression dynamics, with different subunits and isoforms showing distinct profiles: *HIF1A* (HIF-1α) and *ARNT* (HIF-1β) are CB-enriched, predominantly at fetal/peri-natal stages, while *HIF3A* (HIF-3α) and *ARNT2* (an alternative isoform of HIF-1β) are enriched in the fetal SCG. *EPAS1* (HIF-2α) has the most striking expression pattern, being the most highly expressed transcription factor across all sheep CB samples and one of the most abundant transcripts overall in cluster C3 (Table 3). This parallels the data from mice, which also identified *HIF-2α/EPAS1* as one of the most abundant transcripts in mouse CB (Zhou *et al*., 2016; Gao *et al*., 2017; Prange-Barczynska *et al*., 2023). However, our data also reveals that *HIF-2α* expression increases with CB maturation, in particular manifesting a step increase post-natally (Figure 4C-f, Figure 10-C3).

The next most highly expressed transcription factor in C3 is *INSM1* (Insulinoma 1); one of the most highly enriched genes in the adult CB that also shows developmentally increasing expression (Table 1A, Table 3, Figure 10A). INSM1 is highly expressed in many neuroendocrine tissues and used as a marker of neuroendocrine tumours (Möller *et al*., 2024) . We confirmed high INSM1 expression at the protein level in the sheep CB (Figure 10B), showing that it behaves as a CB marker in this species. Like *HIF-2*α*, INSM1* manifests a marked increase in expression post-natally. Other C3 genes encoding transcription factors, such as *TCF4* and *RORA*, show a more progressive increase with development, so may regulate different functions. Interestingly, *MYCN* shows a striking peak in expression at post-natal day 15 but is otherwise expressed at relatively low levels (Figure 10A-C3).

Some transcription factors in other clusters also show progressive increases in CB expression with more complex patterns of differential expression between tissues. *PHOX2B* is overall more highly expressed in the SCG but shows a greater increase in expression over time in the CB, peaking in adulthood (fig 10A -S2). A similar pattern is seen with *PHOX2A*, although this remains most highly expressed in the SCG (Figure 10A-S3). *PHOX2A/B* are important for autonomic nervous system development, along with other transcription factor genes in SCG-enriched clusters, including those known to mediate key stages of sympathoadrenal development. These mostly peak at fetal stages (e.g. *SOX10, ISL1, GATA2, GATA3* (S1)) but some peak post-natally (e.g. *HAND1* (S3)).

Transcription factors with CB-enriched expression at fetal stages include others implicated in developmental processes, such as regulation of retinoic acid signalling (*NR2F2, RARA*) (Figure 10A-C1), and organ maturation via thyroid hormone signalling (*THRB*) (Figure 10A-C2) and glucocorticoid signalling (*NR3C1* – glucocorticoid receptor) (Figure 10A-C1). In addition to the fetal peak in *NR3C1* expression, gene set enrichment analysis shows progressive CB enrichment for glucocorticoid target genes, which becomes significant by adult stages (Figure 11).

**Figure 11.**
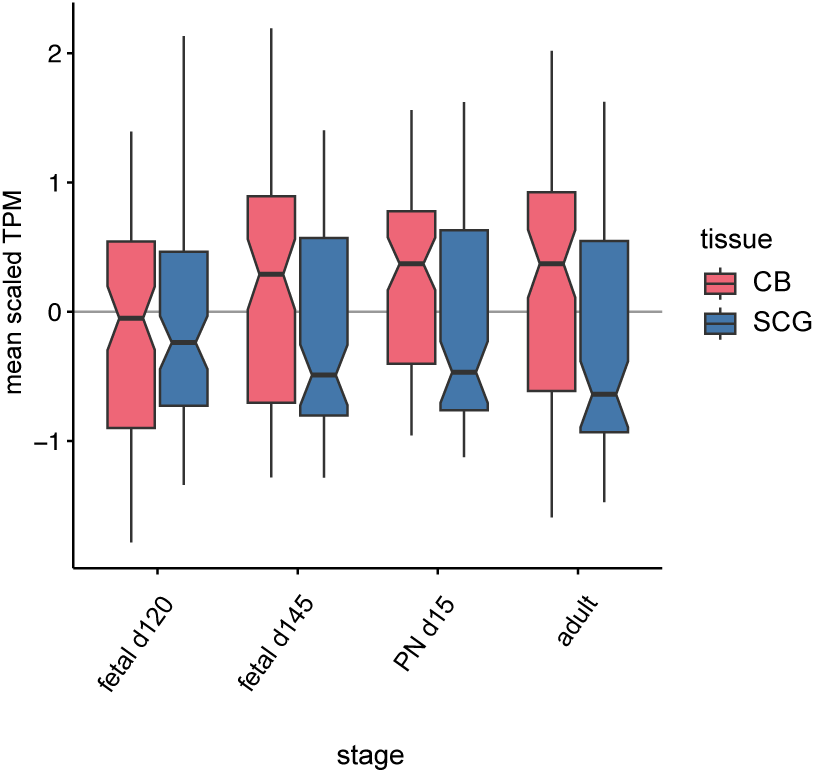
Glucocorticoid receptor target gene expression. Gene set enrichment analysis for glucocorticoid receptor/ response element target genes in CB vs SCG samples at each time point.

## Discussion

We present an extensive, in-depth data-set on CB gene expression, incorporating the first description of the CB transcriptome in fetal life, and the first CB RNA sequencing analysis in a non-rodent species. We find many features of the CB gene expression signature to be conserved across species and already established in fetal life from the earliest stages when oxygen-sensitive chemoreflexes can be reliably measured. By examining how expression of these genes changes with development we have further refined a set of CB-enriched genes that correlate with physiological changes in chemosensitivity. At the core of this gene expression profile, the most over-expressed genes encode variant isoforms of HIFα subunits (*EPAS1*/HIF-2α), regulators of complex IV (*NDUFA4L2, HIGD1C*), and regulators of cell signalling (*RGS5, DGKH*).

It is worth noting that these exceptional levels of transcript expression may correlate with functional significance, but also mechanisms of regulation. HIF-2α is subject to oxygen-dependent degradation, regulated by the oxygen-sensitive PHD enzymes (Kaelin & Ratcliffe, 2008), so the high *EPAS1* transcript expression may reflect high turnover enabling rapid, dynamic regulation of HIF-2-dependent responses. This may be true of other oxygen-labile products, such as RGS5, which is also one of the most abundant transcripts in CB samples. RGS5 is regulated by the oxygen sensitive cysteine dioxygenase ADO and hence targeted for oxygen-dependent degradation via the N-degron pathway (Masson *et al*., 2019), so may also be subject to rapid turnover.

Mechanisms have been proposed for how HIF-2 and the variant isoforms of complex IV could contribute to oxygen chemosensitivity, supported by some experimental data, and our gene expression data further enhances this evidence. Our data also emphasizes the striking CB-enriched expression of certain signalling secondary messengers, particularly *RGS5* and *DGKH*, which are not just highly expressed but increase in parallel with gains in chemosensitivity. Although the significance of these signalling pathways in CB cells has yet to be defined, secondary messenger molecules are of interest as they may be well placed to link metabolic sensing to events at the plasma membrane. Of note, diacylglycerol (DAG) has been shown to directly inhibit TASK-1 and TASK-3 conductance, with similar effects produced by DAG analogues and inhibition of DAG kinases (Wilke *et al*., 2014). DGK activity requires ATP, so it is possible, depending on the dynamics of ATP binding, that this could be sensitive to metabolic flux and facilitate hypoxic signal transduction. For example, if ATP binding by DGKH is sufficiently weak, then acutely reduced cellular ATP could impair DGKH activity, allowing accumulation of DAG that inhibits TASK channels to cause membrane depolarisation. We also find similar CB-enriched gene expression profiles for enzymes catalysing the generation DAG, including phosphatidic acid phosphatases, which could support a high turnover and rapidly responsive changes in DAG levels. Such a mechanism would provide a specific link between the oxygen-sensitive mitochondrial and membrane responses characteristic of CB type I cells. It would also be consistent with the extensive metabolic phenotype we observe at the gene expression level, which would be predicted to result in a low basal metabolic rate, low oxygen consumption, and low flux of ATP/GTP, potentially facilitating the signalling of oxygen levels independently of changes in metabolic demand.

A further aim of this work was to identify factors regulating the development of CB cells and maturation of chemosensory function. These could be used to develop protocols for the differentiation of oxygen chemosensory cells *in vitro,* for instance, from embryonic or induced pluripotent stem cells, or other permissive cell types. We identify a diverse range of CB gene expression characteristics that provide insights into the developmental cues and maturation factors these cells are likely to experience. One of the most consistent features is high relative expression of genes for the organisation of the extracellular matrix. This could have a mechanical role, maintaining CB tissue structure in the pulsatile, peri-arterial niche. It may also provide biochemical signals either directly, for example through the hippo signalling pathway which responds to mechanical cues and is CB-enriched (Figure 6-C1), or indirectly through effects on growth factor signalling. Features of CB transcription factor profiles indicate other potential signals, with nuclear receptors for ligands including retinoic acid and vitamin D3, which may be important for directing differentiation. Other pathways may be more cell intrinsic, for example, circadian rhythms exert variation on ventilatory control and chemoreflexes, and we find several CB-enriched transcription factors with circadian clock functions (*RORA, BHLHE41*-C3, *NR1D1-* cluster C2 (Figure 10)).

Our focus on transcription factors that show progressive, high CB expression (mostly cluster C3) identifies several factors with plausible roles in CB development: INSM1, while not previously studied in the CB, is necessary for normal development of adrenal chromaffin cells (Wildner *et al*., 2008). These adrenal cells have some functional features and ontological origins in common with CB type I cells, including evidence of oxygen chemosensitivity during fetal/neonatal life. TCF4 is mutated in Pitt Hopkins syndrome, causing clinical features including abnormalities of ventilatory control and apnoea (Amiel *et al*., 2007; Zweier *et al*., 2007), which could reflect abnormalities of CB chemoreceptor development. Together with previous work showing that *HIF-2α* is necessary for perinatal survival of CB type I cells (Macias *et al*., 2018), this suggests that the C3 cluster is likely to be enriched for transcription factors regulating the development and/or maintenance of chemosensitive CB cells. PHOX2B is essential for normal development of ventilatory control pathways, with human mutations causing congenital hypoventilation syndrome (Ondine’s curse). It regulates development of both the SCG and CB, with genetic *Phox2b* knock-out causing agenesis of the SCG and involution of the CB in late gestation (Dauger *et al*., 2003). We find that *PHOX2B*, while predominantly SCG-enriched, shows a marked increase in expression in the post-natal and adult CB, raising the possibility that it contributes to both organogenesis and late CB survival and maturation.

In normal peri-natal physiology, a late gestation surge in fetal plasma glucocorticoids mediates maturation of multiple organ systems, for example, lung surfactant production, and antenatal glucocorticoid therapy is used clinically to accelerate these processes prior to threatened pre-term delivery (Fowden *et al*., 1998; Jellyman *et al*., 2020). We identify up-regulation of glucocorticoid receptor (*NR3C1*) expression in the fetal CB, along with progressive enrichment of glucocorticoid target genes with CB maturation. Previous experiments on pregnant sheep have shown that exogenous corticosteroid treatment enhances fetal CB chemoreflexes, such as the fetal brain-sparing circulatory response (Fletcher *et al*., 2003; Jellyman *et al*., 2005). Together with the data reported here, this suggests that the physiological glucocorticoid surge in the fetal circulation contributes to normal CB functional maturation towards term. Interestingly, we also observe that the thyroid hormone receptor (*THRB*) is up-regulated in the CB during the peri-natal period. While this has not been tested as a modulator of CB function, it has established roles in the maturation of other organ systems (Chattergoon *et al*., 2012) and could serve to enhance perinatal chemoreflexes.

We also sought to understand the factors that contribute to CB dysfunction in pathologies including neonatal apnoea and sudden infant death syndrome (SIDS). SIDS is a heterogenous syndrome comprising multiple aetiologies, but most cases remain unexplained. Genetic association studies have identified likely heritable cardiac and metabolic causes in approximately 20-30% of SIDS cases (Hertz *et al*., 2016; Neubauer *et al*., 2017) building the case for post-mortem genetic testing. Ventilatory control has been a focus of SIDS research, with abnormalities of ventilatory patterning (Schechtman *et al*., 1996), apnoea (Kahn *et al*., 1992) and CB morphology observed in SIDS cases (reviewed in Porzionato *et al*., 2018). Focussed screening of genome wide exome data from a cohort of 155 SIDS cases (Neubauer *et al*., 2017) specifically examined 11 genes implicated in central and peripheral chemoreception. In 3% cases, this identified potential pathogenic variants in chemoreceptor genes, including *KCNJ16* (inward rectifying potassium channel 5.1), *KCNMA* (maxi-K channel), *OR51E2* (olfactory receptor) and *PHOX2B* (Neubauer *et al*., 2022), which could impact peripheral and/or central chemoreception. This yield from such a small gene panel, including only 5 genes implicated in CB function, suggests that genetic abnormalities of ventilatory control in SIDS merits further study. Expanding the chemoreceptor gene list to include those we find to correlate with chemoreceptor maturation could enhance the genetic analysis of SIDS and further develop insights into SIDS aetiology and ventilatory control. Interestingly, SIDS-related genes were identified in gene ontology of the S1 (early SCG-enriched) cluster, which may reflect the broader importance of the autonomic nervous system in SIDS pathophysiology.

Going forward, we envisage that this dataset will provide a valuable resource for generating and evaluating hypotheses on carotid body function and development, with respect to normal physiology and pathology, and to inform the rational design of stem cell differentiation protocols and cellular models of oxygen chemosensitivity.

## Methods

### Ethics statement

All procedures were performed under the Home Office Project Licence PC6CEFE59 under the Animals (Scientific Procedures) Act 1986 Amendment Regulations 2012, following ethical review by the University of Cambridge Animal Welfare and Ethical Review Board (AWERB).

### Post-mortem tissue sampling

Ewes and their fetuses (gestational age day 120 or day 145) were humanely killed by overdose of sodium pentobarbitone (0.4 ml.kg^−1^ IV Pentoject; Animal Ltd, York, UK), administered by injection of the maternal jugular vein and the fetus exteriorized by Caesarean section. The umbilical vessels were cut and fetal dissection commenced after confirmation of death. Post-natal lambs similarly underwent euthanasia with a lethal overdose of sodium pentobarbital (200 ml/kg IV Pentoject; Animalcare Ltd., York, UK) administered by injection of the jugular vein. Adult sheep (age ∼10-12 months) were sourced from an abattoir, where the animals were slaughtered by the standard method and the heads immediately removed for dissection. Neck dissection and identification of the CB location at the occipital branch of the carotid artery was performed using previously described approaches (Appleton & Waites, 1957). The origin of the occipital artery, together with the carotid sinus nerve, ganglioglomerular nerve and SCG were removed and transferred on ice cold PBS to a dissecting microscope, where the CB and SCG were identified and dissected free of surrounding tissue. These samples were snap frozen in liquid nitrogen for later processing.

### Sample preparation and RNA sequencing

The frozen samples were lysed in RNA lysis buffer (Qiagen) and refined to a fine suspension using a hand-held motorised homogeniser with a 5 mm probe (Pro-Scientific). RNA extraction was performed using the RNA Clean & Concentrator-5 kit (Zymo) and tested by Bioanalyser, with all samples used in the final analysis having RIN values of 6.1-8.0.

Libraries were prepared using the Kapa RNA hyperprep kit with Riboerase (Roche) and sequenced by NovaSeq with paired end read lengths 100 and 30-50 million reads per sample.

We did not independently analyse gene expression by sex due to the number of pregnancies that would be required to match male and female subjects at each time point. In each case we selected samples from 2 female and one male, with the exception of fetal day 145 which used two males and one female. 3 biological replicates were used per time point, although at the time of analysis one sample (fetal day 145 SCG) was excluded.

### Code availability

The scripts that were used to analyse the RNA-Seq data are available on GitHub (https://github.com/YoichiroSugimoto/20250110_Sheep_CB).

### RNA-Seq data analysis

From the RNA-Seq data, the transcript abundance was estimated using Salmon (version 1.4.0)(Patro *et al*., 2017) with the following parameters: --validateMappings, --seqBias, -- gcBias, and --posBias. The reference transcript data of Ovis aries (sheep) genome assembly: ARS-UI_Ramb_v2.0 (NCBI RefSeq assembly GCF_016772045.1) were used.

For the principal component analysis, the transcript abundances were transformed using the variant stabilisation transformation function of DESeq2 (version 1.42.1) (Love *et al*., 2014) . The top 500 genes in terms of the variance of the transformed values between samples were incorporated in the analysis. The DESeq function of DESeq2 was used to calculate the fold change, the moderated fold change, and statistical significance of the differential gene expression between the two conditions using the Wald test (Zhu *et al*., 2018) .

To define the functional class of sheep genes, orthologous genes were mapped between sheep and human genes using the g:Orth function of g:Profiler (Kolberg *et al*., 2023). The classification of human gene orthologs, which are better annotated, were used in gene ontology analysis. Further analysis of molecular pathways was performed using the Gene Ontology consortium GO Enrichment Analysis tool (Ashburner *et al*., 2000; Thomas *et al*., 2022) (version 10.5281/zenodo.15066566, release date 2025-03-16). Identification of genes encoding transcription factors was performed using the functional annotations on the PANTHER database (PANTHER19.0). For Fig. 9A, functional class of genes involved in metabolic pathways was defined by KEGG orthology, as described previously (Sugimoto & Ratcliffe, 2022) . Volcano plots (Fig. 9B) were plotted using gene lists for Mitochondrial respiratory chain complexes (HGNC gene group 639), Glycolysis (PANTHER accession P00024), and the TCA cycle (KEGG M3985). Glucocorticoid receptor target genes were defined by the Human Gene Set of GRE_C from the Molecular Signature Database (Subramanian *et al*., 2005) .

Hierarchical clustering of genes was performed for genes that showed significant (FDR < 0.1 and absolute value of moderated log2 fold change > 1) differential expression between CB and SCG samples at any time point. The heatmap illustrates the Z-score of mRNA abundance (normalized as transcripts per million) of each gene relative to all replicates, tissues and time points. For clustering, the distance between genes was calculated using Spearman’s rank correlation and the Wald criterion was used as the clustering method. The number of gene cluster groups was determined by minimizing the Kelley-Gardner-Sutcliffe penalty calculated using the maptree package. A csv file of gene expression levels (tpm) and cluster ID for each gene (where applicable) is available here 10.25418/crick.29958284.

### Immunohistochemistry

Carotid bodies and super cervical sympathetic ganglia were fixed overnight in 10% neutral buffered formalin then washed in 70% ethanol prior to processing for paraffin embedding. Paraffin blocks were cut in 4 uM sections, heated to 60° for 1h, pre-treated with hydrogen peroxide block, then boiled in target retrieval solution (citrate, pH 6, Dako/Agilent). Sections were blocked with horse serum then incubated with primary antibody overnight at 4°C (TH (NB300-109, Novus Biologicals), Chromogranin A (SP-1, Immunostar), DGKH (13873-1-AP, Proteintech/ThermoFisher). The sections were washed, then incubated with species-appropriate HRP-coupled secondary IgG polymer (Vector Immpress MP-7401/7402) prior to signal detection with DAB subtrate (Vector laboratories) and mounting in DPX (Sigma-Aldrich). For Masson Trichrome staining, sections were fixed in Bouin’s solution, washed in water, then sequentially stained with Weigert’s iron hematoxylin, Biebrich scarlet-acid fuchsin, phosphomolybdic/phosphotungstic acid, and aniline blue. Stained sections were imaged using the 20x/0.8 UPLXAPO objective on an Olympus Slideview VS200 in a standard configuration, and the images analysed and processed using OMERO (online microscopy environment (Allan *et al*., 2012)).

## Supporting information

summary of gene expression values and cluster by gene

## Author contributions

The study was conceived and designed by were designed by EJH, YS, JAN, DAG, and PJR. EJH, RSM, KJBL, SGF, and NN performed experiments for data acquisition. Figures were prepared and statistical analyses performed by EJH and YS with input from other authors.

The manuscript was prepared by EJH, YS, and PJR, and critically reviewed by all authors for important intellectual content.

## Acknowledgements

Funding for the work was received from the Oxford Branch of Ludwig Cancer Research, the Wellcome Trust (106241/Z/14/Z, 226354/Z/22/Z), MRC (MR/V03362X/1), and the British Heart Foundation (RG/17/8/32924, FS/PhD/25/29633). This work was also supported by the Cambridge Experimental Medicine Initiative, and the Francis Crick Institute, which receives its core funding from Cancer Research UK (FC001501), the UK Medical Research Council (FC001501), and the Wellcome Trust (FC001501).

